# Multi-task representations in human cortex transform along a sensory-to-motor hierarchy

**DOI:** 10.1101/2021.11.29.470432

**Authors:** Takuya Ito, John D. Murray

## Abstract

Human cognition recruits diverse neural processes, yet the organizing computational and functional architectures remain unclear. Here, we characterized the geometry and topography of multi-task representations across human cortex using functional MRI during 26 cognitive tasks in the same subjects. We measured the representational similarity across tasks within a region, and the alignment of representations between regions. We found a cortical topography of representational alignment following a hierarchical sensory-association-motor gradient, revealing compression-then-expansion of multi-task dimensionality along this gradient. To investigate computational principles of multi-task representations, we trained multi-layer neural network models to transform empirical visual to motor representations. Compression-then-expansion organization in models emerged exclusively in a training regime where internal representations are highly optimized for sensory-to-motor transformation, and not under generic signal propagation. This regime produces hierarchically structured representations similar to empirical cortical patterns. Together, these results reveal computational principles that organize multi-task representations across human cortex to support flexible cognition.

## Introduction

Humans can perform a variety of tasks in daily life that involve diverse cognitive functions. What are the core neural and computational architectures that facilitate flexible cognition? Current efforts to uncover the neural bases of human cognition typically design carefully controlled experimental paradigms that target specific cognitive functions while measuring spatial patterns of brain activity^1,2^. While this approach has been fruitful for identifying *where* cognitive processes map to in the brain, it is unable to reveal *how* information is represented within brain regions. In contrast, advancements in data analysis have enabled the characterization of fine-grained representations of sensory stimuli within brain regions of individual subjects^3,4^. However, an overarching understanding of how fine-grained representations are organized across diverse cognitive functions and across brain areas remains poorly studied.

Studies in experimental neuroscience and human functional magnetic resonance imaging (fMRI) have revealed the spatial organization of cognitive functions and specialization across cortex^5^. For example, by mapping the stimulus-response properties of different brain areas, researchers have identified regional correlates of working memory^6^, visual processing^7^, and motor function^8^. More recently, meta-analytic human neuroimaging studies have made progress in characterizing the organization of functional specialization and flexibility by aggregating datasets from many tasks and studies^9,10^, finding functional specificity in unimodal (sensorimotor) areas and flexibility in transmodal (association) areas^11,12^. These prior approaches typically used datasets with brain activations collected from only a single task for each subject. The single-task nature of prior datasets, including meta-analyses which aggregate over them, fundamentally limits their ability to reveal the structure of neural representations within brain regions because fine-grained spatial patterns do not align well across subjects^13^. Thus, studying the detailed representational cortical topography for diverse cognitive processes requires datasets in which multiple tasks are collected per subject, enabling fine-grained representational analyses not feasible with coarse-grained meta-analytic approaches.

The measurement of many tasks from the same subjects would allow for the characterization of multi-task representational structure – a key element for flexible cognition. Analytically, a leading approach to facilitate this endeavor is representational similarity analysis (RSA)^3,4^. RSA measures geometrical properties of representations within a brain region by comparing the similarity of multivariate activations (e.g. across voxels in fMRI) across different task conditions. Representational geometry then can be compared across brain regions^4^, as well as between brain data and computational models^14,15^. While this approach can identify task-relevant representational geometries for specific brain regions, previous studies have typically been limited to isolated tasks in specific domains (e.g., perceptual tasks)^16,17^. This limits the interpretation of representational geometry within and between brain regions, since such tasks only recruit a small subset of the diverse cognitive processes of which humans are capable.

Representational geometry provides a unifying framework for relating neural data and computational models to understand principles of perceptual and cognitive function. In particular, studies have compared representations from different brain regions in human fMRI to the internal representations in deep neural network models of sensory processing, including in vision and audition^15,18^. Despite active interest in modeling multi-task cognition^19,20^, models of multi-task cognitive function have not been related to functional differences across brain regions due in part to lack of empirical characterization of multi-task representations in the brain.

Here we use a human fMRI dataset with 26 tasks per subject to probe the computational properties of multi-task representations across the entire cortex. We first characterized the multi-task representational geometry within each cortical area. Analysis of how these representations align and vary across areas revealed a hierarchical organization of representational variation, which spanned from sensory, to association, to motor areas. We also quantified the dimensionality of multi-task representations for each cortical region. Along this sensory-association-motor axis, we found that multi-task representational dimensionality first compressed from sensory to association areas, and then expanded from association to motor areas. To investigate computational principles of multi-task representations, we trained multi-layer, feedforward artificial neural networks to transform empirical visual to motor representations. Compression-then-expansion organization in network models emerged exclusively in a training regime where internal representations are highly optimized for sensory-to-motor transformation, and not under generic signal propagation^21,22^. Moreover, this regime produces hierarchically structured representations similar to those in the brain. Together, our findings reveal the hierarchical organization of multi-task representations in the human brain, and establish a framework to develop neurally-inspired computational architectures for multi-task cognition.

## Results

### Analytic approach to studying multi-task representations

RSA of empirical data was central to our data analytic approach (Fig. 1)^4^. RSA approximates the representational geometry of a set of multivariate vertex activations by comparing the similarity of activation vectors across different task conditions. By performing RSA on each brain region separately, we could produce representational similarity matrices (RSMs) for every brain region (Fig. 1a). Critically, RSMs were calculated at the individual-subject level, which ensures that fine-grained representational geometries would not be lost through cross-subject averaging of activations. Because each subject’s cortex is registered to a common atlas and parcellated into discrete areas^23^, individual-level results can be combined to characterize the average RSM for each cortical area. With RSMs from all cortical areas in the Glasser parcellation, we performed a variety of new analyses. First, we compared the similarity of RSMs between cortical areas — i.e., the representational alignment (RA) — and mapped the topography of RA variation across cortex (Fig. 1b). Second, we characterized properties of representational geometry, such as dimensionality, and how they vary across the cortical hierarchy (Fig. 1c). Finally, we built computational models of representational transformation across the cortical hierarchy and evaluated the correspondence between the model’s internal representations and empirical observations (Fig. 1d).

**Figure 1.**
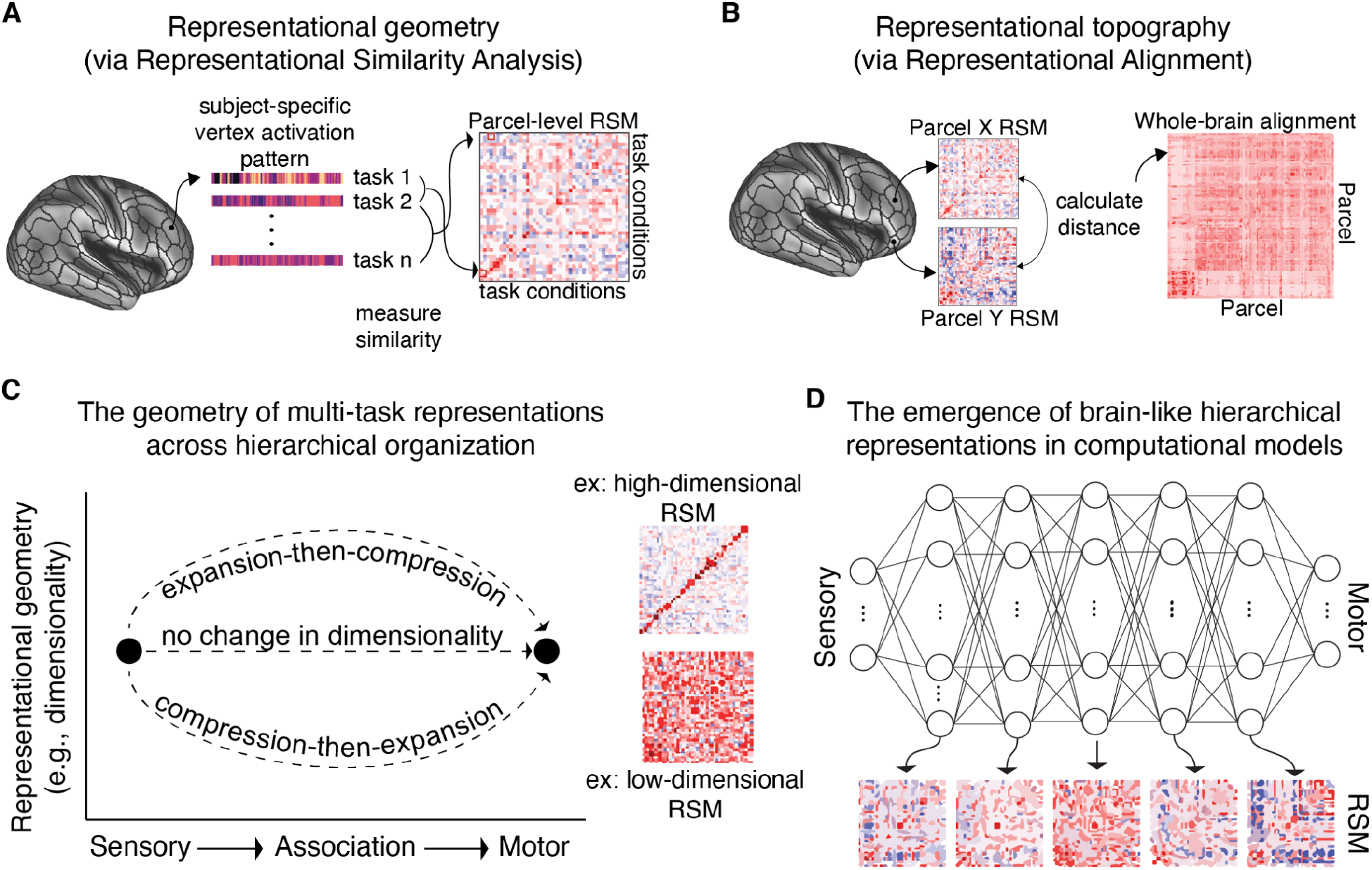
Overview of analytic approaches to study the geometry and topography of multi-task representations in fMRI data. **a)** Representational geometry of brain parcels is characterized by RSA applied to subject-specific vertex activation patterns within each parcel^4^. Using subject-specific activation patterns ensures that fine-grained representational geometries would not be lost through cross-subject averaging. This enables estimation of a representational similarity matrix (RSM) for each brain region using vertices within that region. **b)** Using each region’s RSM, we can characterize the topography of representations by measuring the representational alignment (RA) – the similarity of regional RSMs – between all pairs of brain regions. **c)** We next ask how representational geometry changes across the sensory-motor hierarchy. An example of a high-dimensional geometry is one with a strong diagonal, but weak off-diagonal. In contrast, a low-dimensional geometry is one with a lack of structure in the RSM and high similarity in activation patterns between conditions. **d)** Given the empirical results, we identify the conditions by which similar hierarchical representations emerge in the internal layers of feedforward neural network models trained to produce sensory-to-motor transformations.

### A publicly available dataset with 26 cognitive tasks

Characterizing fine-grained multi-task representations across cortex requires a dataset that collected many tasks per subject, which is rare in neuroimaging. We used the multi-domain task battery (MDTB) human fMRI dataset with 26 cognitive tasks, comprising up to 45 unique task conditions, collected per subject^24^ (Fig. 2a). Data were collected across four imaging sessions, enabling within-subject cross-validation analyses across task conditions. This facilitated the characterization of multi-task representational geometries at the within-subject level across the entire cortex (Fig. 2c).

**Figure 2.**
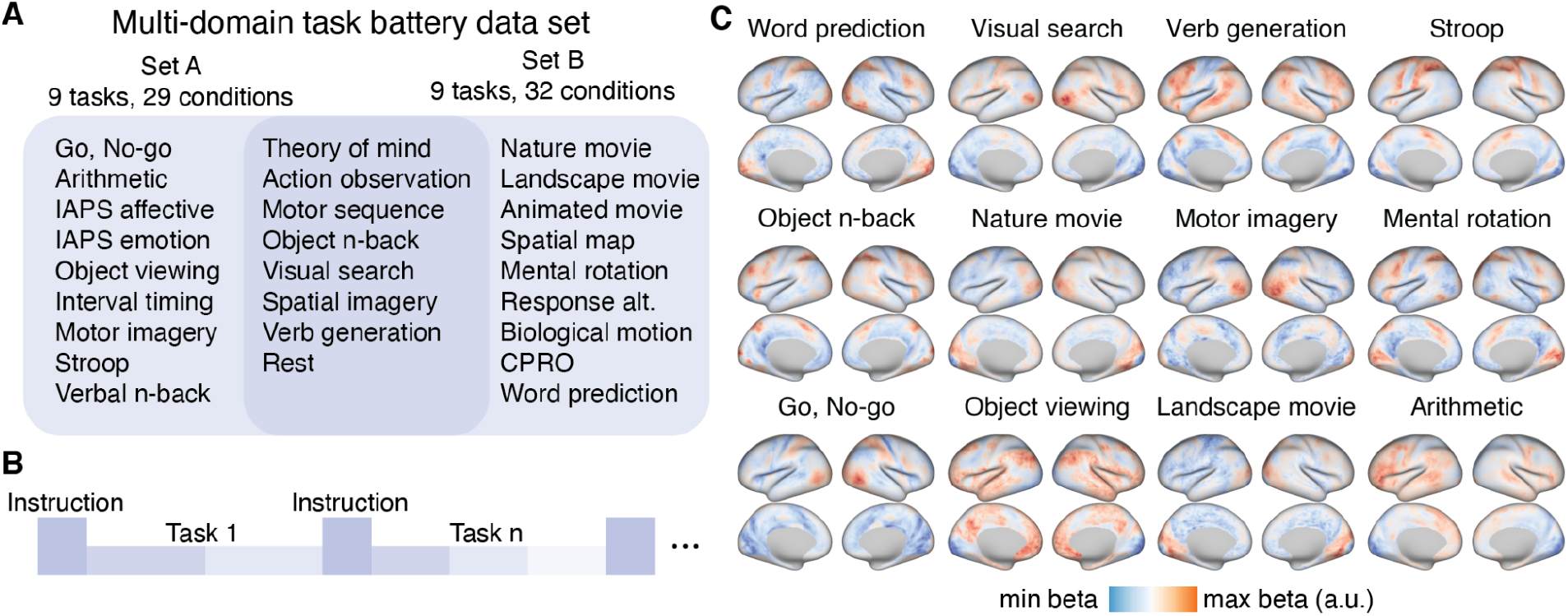
Leveraging the multi-domain task battery (MDTB) dataset to investigate multi-task representational geometries. **a)** The MDTB dataset consists of 26 distinct tasks with up to 45 unique task conditions, and was previously made publicly available^24^. The tasks were split across two sets. Every subject performed each set of tasks twice across different fMRI sessions. **b)** Task blocks were interleaved across each fMRI session. For each block, instructions were presented for 5 seconds, followed by a task which was performed continuously for 30 seconds until the subsequent block. **c)** Whole-cortex group-level activation maps for 12 of 26 cognitive tasks. (See Supplementary Fig. 1 for all task activation maps.)

### Measuring the similarity of task content between brain regions with RA

We used the multi-task representational geometry of each cortical area to characterize the topography of representational variation across cortex (Fig. 3a). The representational geometry of each area was reflected in its RSM. The RSM of each cortical area was computed using the vertices within each of 360 areas in the multi-modal parcellation of Glasser et al. atlas^23^. We specifically chose this parcellation due to improved delineation of somatotopic and visuotopic areal organization (motor and visual receptive fields, respectively) that are not accounted for in atlases defined solely on resting-state fMRI^23^. To characterize the inter-regional similarity of multi-task representations, RA was computed as the cosine similarity of the RSMs between two regions (Fig. 3b), which produces a whole-cortex RA matrix (Fig. 3c). One common way to characterize the large-scale organization of cortex is through resting-state functional connectivity (RSFC)^25,26^. We found that the similarity of multi-task RA and resting-state was moderate (ρ = 0.37, p < 0.0001; Fig. 3d), suggesting that the cortical multi-task RA matrix offered unique information from typical RSFC analyses.

**Figure 3.**
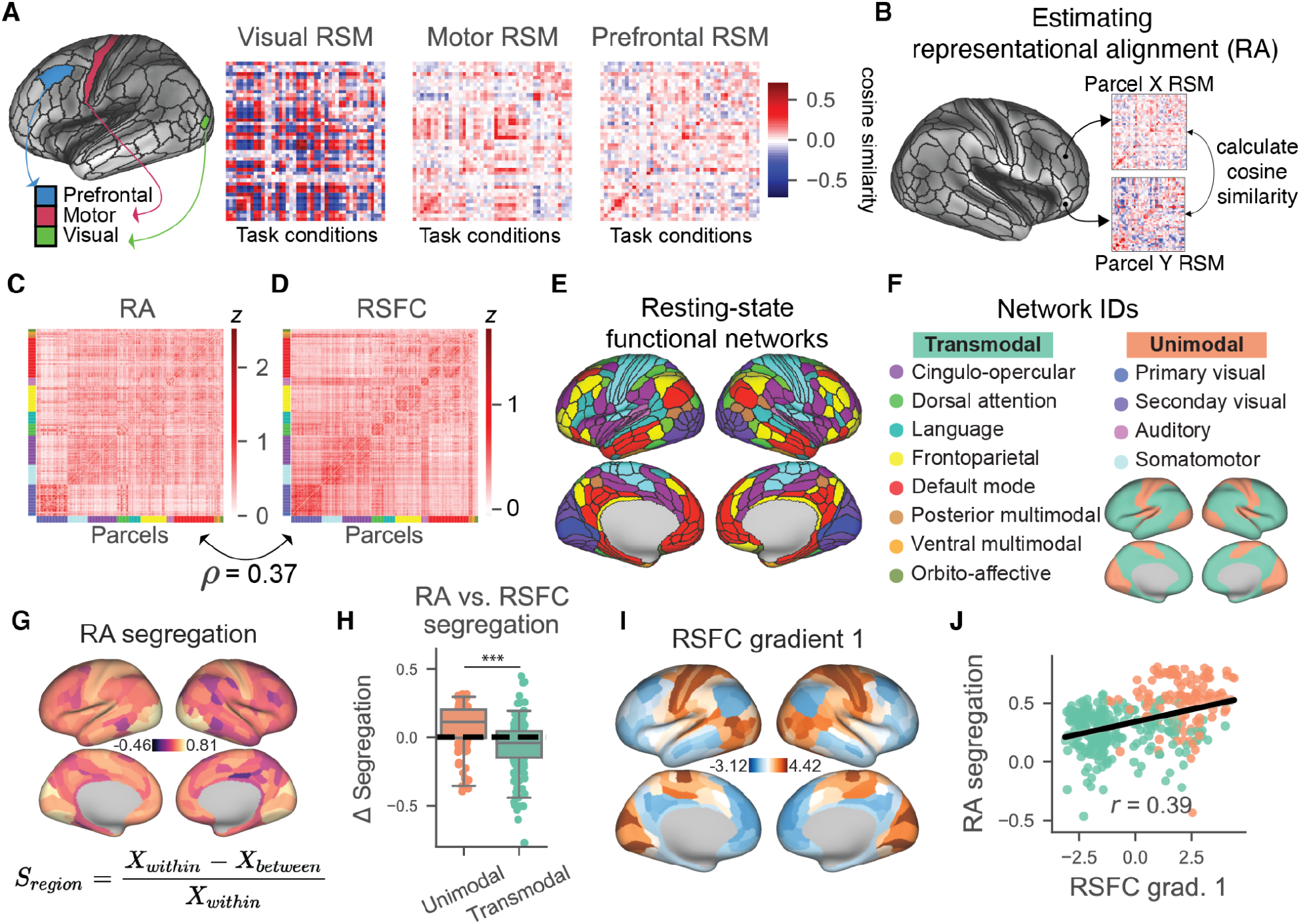
The cortical organization of multi-task representations. **a)** Three example RSMs taken from visual, motor, and prefrontal areas. RSMs consisted of 45 task conditions, cross-validated across imaging runs. **b)** RA between pairs of cortical areas is quantified by measuring the cosine similarity between their RSMs. **c)** RA and **d)** RSFC for all pairs of cortical areas. **e, f)** Previously identified RSFC networks. **g)** Segregation (*S_region_*) of the RA and RSFC matrices, defined as the difference of within-network (*X_within_*) and between-network (*X_between_*) values, divided by within-network values. **h)** Unimodal regions have greater segregation for RA relative to RSFC, and transmodal regions have less segregation for RA relative to RSFC. This was despite no difference in overall segregation between RA and RSFC. **i, j)** The cortical topography of RA segregation is correlated with the RSFC principal gradient, a proxy of intrinsic hierarchy. (Green/red dots reflect transmodal/unimodal regions, respectively.) (***p=<0.001).

### Network topography of RA relates to the brain’s intrinsic organization

We next characterized RA in the context of the well-established functional network organization of the brain. RSFC can be used to assign each cortical area to a functional network^25–27^, and this network organization is strongly related to cognitive task activation patterns^28^. We measured the segregation of RA in relation to the segregation of RSFC. Conceptually, segregation measures how isolated a network’s representations/FC are in relation to other networks of the brain. Segregation of a measure (e.g., RA or RSFC) for a given brain region is calculated as the difference between its within-network and between-network values, divided by the within-network value (Fig. 3g,h)^29^. Thus, if a visual region’s RSM was highly local (and unique) to the visual network, then its segregation would be high. First, we found that unimodal regions had significantly higher RA segregation relative to transmodal regions (t_358_=12.99, p<10e-31) (Supplementary Fig. 2b). We did not observe a significant difference in mean segregation between RA and RSFC at the whole-cortex level (t_358_=-1.24, p=0.22). However, we found that RA had exaggerated differences in segregation by network: unimodal networks had significantly higher segregation for RA relative to RSFC (t_112_=-3.33, p=0.001), and transmodal networks had significantly lower segregation for RA (t_244_=-4.24, p<10e-04) (Fig. 3g,h). These results illustrate that representations in unimodal regions were more isolated, while transmodal regions were more distributed.

We sought to evaluate the relationship between RA segregation with previously reported measures of hierarchical organization. Large-scale topographic gradients of cortical organization have been defined using task-free MRI measures^30,31^. Extracting the principal components of the RSFC matrix produces a unimodal-transmodal hierarchical gradient (as the first principal component) and a sensory-to-motor hierarchical gradient (as the second principal gradient)^30^. Moreover, the unimodal-transmodal RSFC gradient has previously been shown to be highly related to the cortical T1w/T2w map (an MRI-contrast correlate of intracortical myelin content)^32^. Thus, we correlated RA segregation with the principal gradient of RSFC (leading component of a principal component analysis on the RSFC matrix; *r* = 0.39, non-parametric p < 0.001; Fig. 3i,j)^30^, and the T1w/T2w myelin map (*r* = 0.36, non-parametric p < 0.001; Supplementary Fig. 2f,g). These findings establish a link between task-free properties of hierarchical organization with multi-task representational topography.

### Task representational dimensionality is associated with the unimodal-transmodal hierarchy

Recent experimental and modeling studies have investigated the role of dimensionality of stimulus representations during task states^33,34^. Representational dimensionality refers to the dimensionality of the task state space. However, most prior studies evaluated the representational dimensionality of either a specific task (e.g., perceptual task) or within a specific brain region, such as the prefrontal cortex^34^. Here we evaluated the representational dimensionality across 45 task conditions, and across the entire human cortex.

We measured the representational dimensionality by measuring the participation ratio of the RSM for each cortical region (Fig. 4a)^35–37^. Intuitively, the participation ratio is a statistical estimate of dimensionality, and is related to the flatness of the eigenvalues of the RSM (e.g., a flatter scree plot corresponds to higher dimensionality). For instance, if the activation pattern is highly similar for all task conditions (a highly correlated RSM), dimensionality will be low. In contrast, if the activation pattern for a task condition is uniquely similar to itself (with low similarity to other task conditions), then dimensionality will be high (Fig. 1c). (Note high-dimensionality corresponds to strong diagonal versus off-diagonal components in the RSM.) In addition, we estimated the multi-task (45-way) decoding as a complementary measure of dimensionality, since decoding has also been previously used to estimate dimensionality^34^.

**Figure 4.**
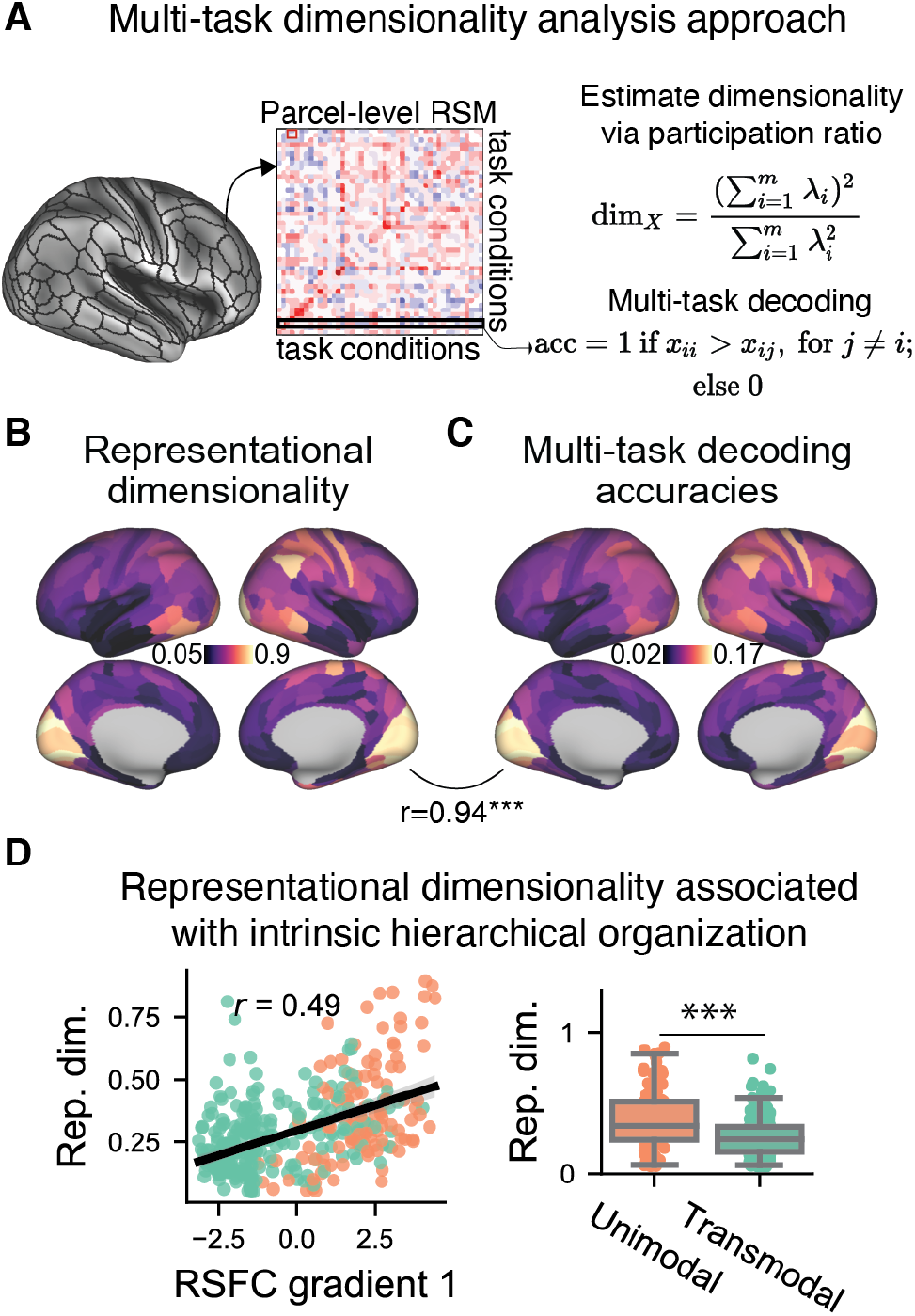
The representational dimensionality of task activations follows hierarchical organization. **a)** We estimated multi-task representational dimensionality using two approaches: 1) Estimating the dimensionality of the cross-validated RSM, where λ_*i*_ refers to the *i*-th eigenvalue of the RSM; 2) Performing within-subject decoding across all possible task conditions (n=45), with cross-validation across sessions (split-half). **b)** The representational dimensionality and **c)** multi-task decoding accuracy of each cortical parcel. **d)** Across cortex, representational dimensionality was positively correlated with the first principal gradient of RSFC, with unimodal regions containing higher representational dimensionality than transmodal regions. Green dots reflect unimodal regions. Red dots reflect transmodal regions. (See Supplementary Fig. 3 for equivalent plots for the decoding approach.) (***p=<0.001)

We found that the representational dimensionality (using participation ratio) and the multi-task decoding accuracy of each region were highly correlated across cortical areas (r=0.94, non-parametric p<0.001) (Fig. 4b,c), suggesting that both approaches were reliable methods for estimating representational dimensionality. Next, we addressed whether representational dimensionality was related to intrinsic hierarchical organization. Indeed, we found that representational dimensionality and multi-task decoding were highly correlated with the RSFC principal gradient (r=0.49, non-parametric p<0.001) and T1w/T2w myelin map (r=0.41, non-parametric p<0.001) (Fig. 4d; Supplementary Fig. 3). These results illustrated that like representational segregation, representational dimensionality was also higher in unimodal regions than in transmodal regions (t_358_ = 6.54, p<10e-9; Fig. 4d). These findings are consistent with previous studies that have found that higher-order association areas typically have relatively low decoding accuracies, even for tasks that heavily involve those regions^38^.

As control analyses, we tested for the effects of parcel size and number of task conditions. Although parcel size and representational dimensionality are positively correlated (r=0.45, non-parametric p<0.001), after conditioning on parcel size as a covariate using linear regression, the associations between representational dimensionality and hierarchical organization remained significant (correlation with RSFC gradient 1, r=0.42, non-parametric p<0.001; correlation with T1w/T2w myelin map, r=0.33, non-parametric p<0.001) (Supplementary Fig. 3c). We tested for robustness to the number of task conditions by randomly sampling subsets of task conditions, and found that the hierarchical differences in representational dimensionality were robust if at least 10 task conditions were included (Supplementary Fig. 4a-c).

### Compression-then-expansion of task representations across the sensory-motor hierarchy

We next sought to map the dominant topographic axis of RA variation across cortex. Thus, we performed a principal component analysis to extract the first principal gradient of cortico-cortical RA. In contrast to the unimodal-transmodal principal gradient exhibited from RSFC (Fig. 3i), RA’s principal gradient exhibited a sensory-to-motor gradient (Fig. 5a). This sensory-motor gradient was similar to the second principal component of RSFC that also reflects a transition from sensory-to-motor cortices (Fig. 5b). Indeed, we found that the sensory-motor RSFC gradient 2 was highly correlated with the RA principal gradient (r=0.59, non-parametric p<0.001), suggesting that in contrast to RSFC, the dominant axis of variation in RA places sensory and motor representations on opposite ends (Fig. 5c).

**Figure 5.**
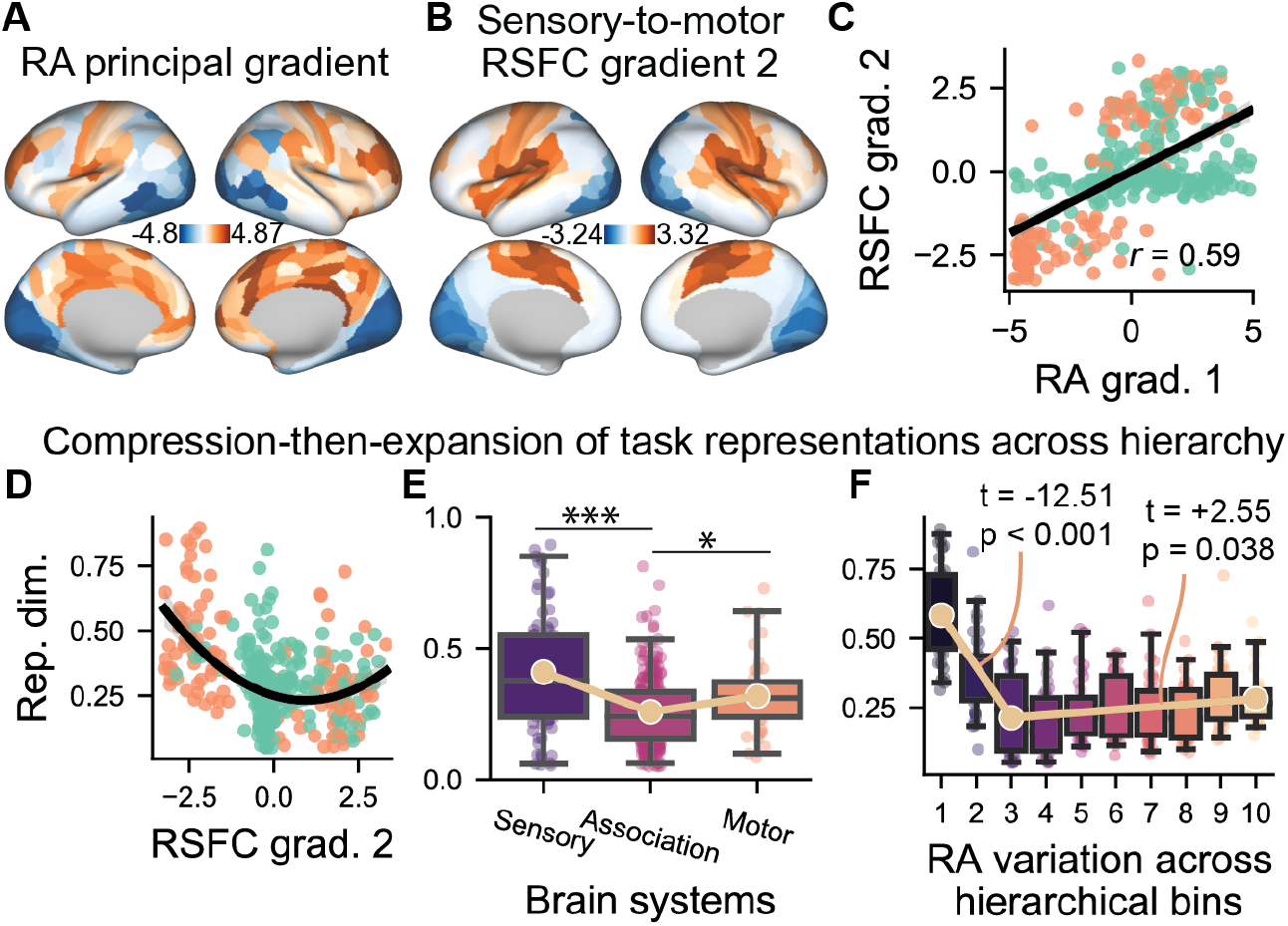
Principal component of the RA matrix reveals a sensory-motor gradient that compresses-then-expands task representations. **a)** Similar to estimating intrinsic RSFC gradients, we extracted the principal component of the cortical RA matrix, which showed striking similarity to the **b)** 2nd RSFC (i.e., sensory-motor) gradient. **c)** Correlation between the sensory-motor RSFC gradient and the principal RA gradient. **d)** We plotted the representational dimensionality against the RSFC sensory-motor gradient, and found that the 2nd-order convex polynomial model was a better fit than a 1st order polynomial model, and an exponential decay model (Supplementary Fig. 5). This suggested that representational dimensionality compressed then expanded across the sensory-motor hierarchy. **e)** Same as in **d**, but after grouping together sensory (visual and auditory network), motor (somatomotor network), and association (all other network) parcels according to network affiliation. **f)** Same as in panel **d**, but after placing regions into 10 bins according to the RA hierarchy (i.e., binning regions together with similar loadings). Using a continuous piecewise linear model, we found a significant negative-then-positive slope best accounted for dimensionality, consistent with compression-then-expansion of dimensionality. (Note statistical tests in panels **e** and **f** are two-tailed tests.) (***=p<0.0001, *=p<0.01)

We next sought to evaluate how representational dimensionality changed across this sensory-motor hierarchy, using the sensory-motor RSFC gradient 2. We fit several competing statistical models to evaluate how representational dimensionality (dependent variable) changed as a function of the sensory-motor gradient (independent variable): linear, quadratic, and exponential decay models. We computed the Bayesian information criterion (BIC) and the Akaike information criterion (AIC) for model selection, which takes into account the maximum likelihood of each model while penalizing models with more free parameters. We found that the quadratic model had the lowest BIC and AIC of all models (Supplementary Fig. 5), suggesting a convex quadratic fit best explained representational dimensionality as a function of sensory-motor hierarchy (Fig. 5d). The quadratic dependence was robust to task condition subsampling (Supplementary Fig. 4d-f). This analysis revealed that representational dimensionality first compressed, then expanded, across the sensory-motor hierarchy.

To verify compression-then-expansion from sensory-to-motor systems, we grouped regions together by their system-level association. We found that both sensory and motor systems had greater dimensionality than association regions (Sensory vs. Association, t_319_ = 7.22, p < 10e-11; Motor vs. Association, t_283_ = 2.59, p = 0.01) (Fig. 5e). Finally, to further establish compression-then-expansion along the RA hierarchy, we created 10 bins of brain regions sorted by the loadings of the RA principal gradient (Fig. 5f). We fitted a continuous piecewise linear regression model, varying the breakpoint between the two line segments at every bin between 2 and 9, and selecting the model with highest R^2^, which resulted in the piecewise model with a breakpoint at bin 3 (see Methods). We then tested the statistical significance for the coefficients of the piecewise regression (i.e., for a negative slope from bin 1 to 3 and a positive slope from bin 3 to 10). Indeed, confirming the compression-then-expansion of representational dimensionality across the sensory-motor hierarchy, we found a negative slope from bin 1 to 3 (t_7_ = −12.51, p < 0.001) and a positive slope from bin 3 to 10 (t_7_ = 2.55, p = 0.038) (two-sided test).

### Compression-then-expansion of task representations in a feedforward ANN emerges during rich training

We next sought to investigate computational mechanisms that could potentially explain the compression-then-expansion of representational dimensionality observed in fMRI data. Interestingly, a recent study of feedforward artificial neural network (ANN) models trained on object recognition tasks observed expansion-then-compression of representational dimensionality, counter to our empirical findings in the brain^39^. Thus, we first asked how the compression-then-expansion phenomena of representational dimensionality emerges in ANNs. We used multilayer, feedforward linear ANNs with tied weights to study how fMRI activations in visual areas were successively transformed into motor activations under different learning regimes. (Corresponding analyses of an ANN without tied weights are shown in Supplementary Fig. 6.)

The representations learned by an ANN can depend strongly on the training regime. Prior research has shown that small alterations to weight initialization parameters can greatly impact the structure of the learned hidden representations in ANNs^21,22^. Specifically, those studies found during a “rich” training regime (in which network initializations had small weight variances), ANNs learned lower-dimensional and structured representations. In contrast, during a “lazy” training regime (in which weight initializations had large variances), task performance was achieved by randomly projecting input features into a high-dimensional embedding in hidden layers. Therefore, we examined dimensionality and representations as a function of the rich and lazy training regimes.

Using the sensory-motor RSFC gradient 2 (Fig. 5b), we selected two brain regions on opposite ends of this axis (defined by the lowest and highest loadings). This resulted in a visual and a motor parcel (Fig. 6a). Note that these sensory and motor parcels had highly similar RSMs to V1 and M1, respectively, suggesting that the gradient-selected parcels were appropriate to model early sensory to late motor transformations in data (Supplementary Fig. 7a-c). We then took the fMRI activations of each of the 45 task conditions, and trained an ANN with 10 hidden layers using weight initializations with different standard deviations (SD) to predict motor fMRI activations (Fig. 6b). We trained 20 random initializations for each specific weight initialization (ranging from 0.2 to 2.0, in 0.2 increments).

**Figure 6.**
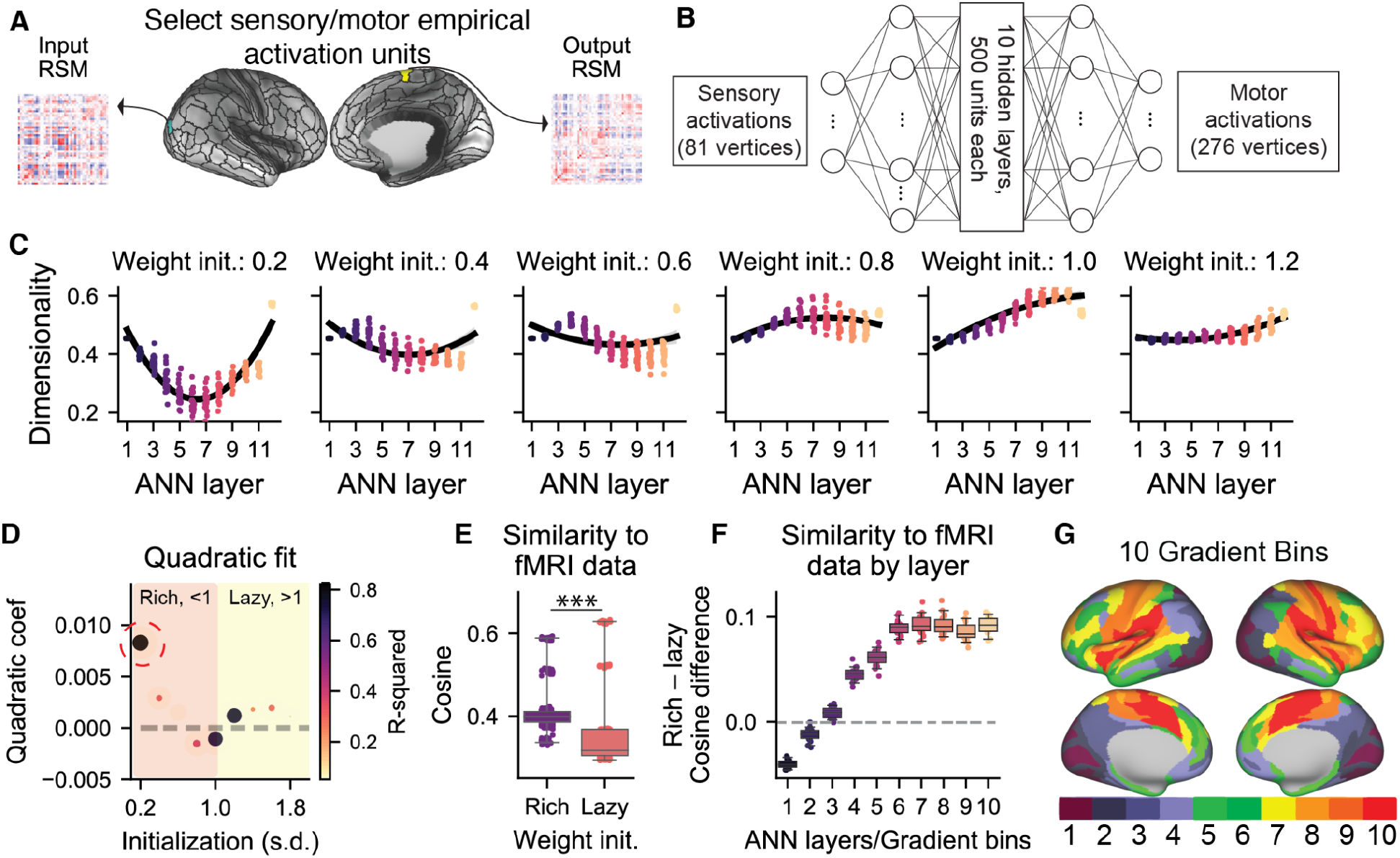
Multi-task representations in human cortex were consistent with ANN representations trained in a rich regime. **a)** We identified the brain parcels at the bottom (visual parcel) and top (motor output parcel) of the sensory-motor RSFC gradient 2, and extracted their vertex-wise task activation patterns. **b)** We trained a feedforward ANN with 10 hidden layers to predict multi-task activations in the motor parcel using vertex-wise activations from the visual parcel. **c)** The representational dimensionality (participation ratio of RSMs) of each layer with different weight initialization SDs. We observed compression-then-expansion of representations in rich training regimes. **d)** We fit a 2nd-order polynomial regression to the dimensionality across weight initializations. We plotted the quadratic coefficient (positive for convex) and the overall R^2^ fit to assess how dimensionality changed across ANN layers. R^2^ peaked during rich training regimes, and was consistent with a convex fit. **e)** We compared the overall similarity of the RSMs of ANNs at each layer with the RSMs for every region in the brain, finding stronger similarity when the ANN was trained in the rich regime (<1.0 SD initialization). **f)** We compared the RSMs of ANNs to the empirical brain RSMs at each bin along the sensory-motor gradient for rich (<1.0) and lazy (>1.0) learning regimes. **g)** Empirical gradient bins were defined by partitioning the sensory-motor RSFC gradient 2 into 10 distinct sets of regions sorted by their gradient loadings.

After ANN training converged, we measured the representational dimensionality of each ANN’s hidden layer. We found that rich training regimes (e.g., weight initialization SD = 0.2) showed compression-then-expansion across layers, consistent with empirical data (Fig. 6c). As with the empirical data, we fit a 2nd-order polynomial regression to model dimensionality as a function of ANN layer depth. We found that in the rich regime, in particular for weight initializations starting at an SD of 0.2, the quadratic fit was convex and had higher R^2^ fit (Fig. 6d). This illustrated that rich regimes were consistent with empirical data in producing hierarchical representations that first compressed, then expanded, across layers.

We next sought to evaluate whether representations learned in the rich regimes were more similar to fMRI representations. While ANNs and fMRI data have different spatial dimensions, direct comparison between ANNs and empirical data could be performed in a task-representational space. Therefore, we compared the RSMs of every cortical parcel from data with the RSM of every ANN layer for rich (i.e., less than 1.0 SD weight initializations) and lazy (i.e., greater than 1.0 SD weight initializations) learning regimes. We found that despite keeping fixed all parameters other than weight initializations, hidden representations learned in the rich regime were more similar with those found in empirical data (rich, cosine=0.42; lazy, cosine=0.37; rich vs. lazy, t_198_=15.28, p<10e-34; Fig. 6e). We then partitioned brain parcels into 10 bins of 36 parcels and sorted them according to their loading relative to the sensory-motor RSFC gradient (Fig. 6g). We correlated the RSMs of each bin with each ANN layer according to depth (e.g., similarity of RSMs for ANN layer *i* with fMRI bin *i*). We found that the later 8 out of 10 bins/layers had higher similarity in the rich learning regime (for 8/10, all FDR-corrected p<10e16). While the first two fMRI bins had greater correspondence with the lazy learning regime, these first two bins primarily consisted of visual cortical areas. Empirically, we found visual areas to contain high-dimensional representations. Since the lazy learning regime typically embeds input features in a high-dimensional space, the higher similarity of the lazily trained ANN with visual regions was unsurprising. Thus, with the exception of early visual areas, we find that the rich learning regime has greater correspondence with fMRI data in terms of both representational dimensionality and content.

### Richly trained ANNs learn hierarchical representational transformations

Having modeled the successive transformation of fMRI activations from visual to motor regions in a feedforward ANN, we next sought to characterize the properties of ANNs that contributed to better correspondence with empirical brain data. First, we characterized the structure of representations that emerged in ANNs trained under different learning regimes. This was done through a similar data analytic strategy as in the empirical data: For each layer, we computed its RSM using each of the 45 activation patterns, and then computed the cosine similarity of that layer’s RSM with the RSMs from all other layers (Fig. 7a). This produced a layer-by-layer RA matrix for different weight initializations (Fig. 7d). We found that ANNs trained in the rich regime learned sequentially structured representations that were consistent with structured representations in empirical data (Fig. 7b-d). In other words, adjacent layers had high RA to each other, but distal layers had low RA to each other. We quantified this by calculating the mean of the ANNs RA matrix for each weight initialization (Fig. 7e) and the dimensionality of the RA matrix (Fig 6f,g). Here, dimensionality was approximated by the amount of variance explained by the first principal component of the RA matrix; the larger the variance explained, the lower the dimensionality. We found that the higher (lazier) the weight initialization, the greater the overall RA across layers and the lower the dimensionality (rich vs. lazy cosine difference=-0.12, t_18_=-124.70, p<10e-28; Rich vs. lazy variance explained by first PC diff=-15.52%, t_18_=-99.10, p<10e-26). In contrast to the rich training regime, the lazy regime had nearly no representational structure for weight initializations with an SD greater than 1.2, suggesting that no representational transformations occurred in the hidden layers, and output activations were transformed in only the last layer (Fig. 7d).

**Figure 7.**
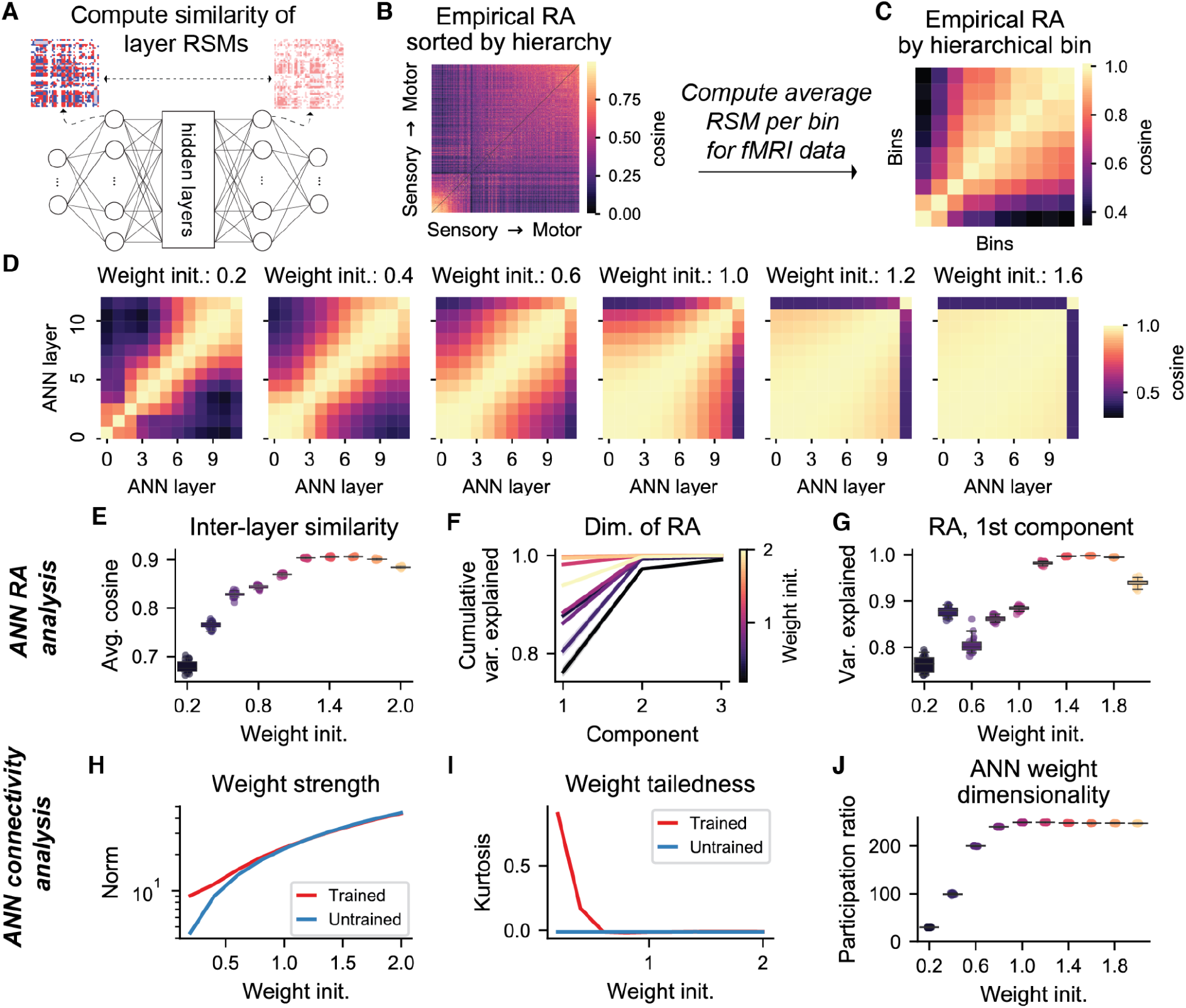
Analysis of the ANN revealed that richly trained ANNs learn diverse and structured representations consistent with empirical data. **a)** We computed the RA between all layers by computing the cosine similarity between the RSMs of each hidden layer. **b)** For comparison, we sorted the empirical fMRI RA by the RSFC sensory-motor (2nd) gradient, and **c)** downsampled it into 10 discrete bins for comparison with the ANN analysis. **d)** The RA for ANNs by layer across weight initializations. ANNs trained in the rich regime (e.g., weight initializations < 1) learn differentiated and structured representations. In contrast, ANNs trained in the lazy regime largely produced impoverished representations that only transformed sensory representations in the final layer. **e)** The average cosine similarity of each RA matrix by weight initialization. **f)** Cumulative variance explained plot of the first three components of the RA matrix under different weight initializations. **g)** Variance explained of only the first principal component of the RA matrix. **h)** Initialized and trained norm of ANN weights as a function of weight initialization. **i)** The kurtosis of the degree distribution during initialization and after training. **j)** Dimensionality of the ANN’s connectivity weights.

Next, we characterized properties of the learned ANN weights. In line with previous work, we first calculated the Frobenius norm of weights under different weight initializations^21^. (Weights with a large Frobenius norm are sampled from a distribution with wide variance.) We found that under the rich training regime, relative weight norms significantly increased from initialization to convergence, relative to the lazy regime (weight norm change, Rich=2.37, Lazy=-0.40, Rich vs. lazy t_18_= 3038.38, p<10e-54) (Fig. 7h). We also calculated the kurtosis of the weight distribution, which measures the tailedness of the weight distribution. Even though we initialized all weights from a Normal distribution (which has a Fisher kurtosis of 0), we found that the kurtosis of richly trained networks increased from initialization, producing a heavy-tailed weight distribution, which is a commonly observed feature of empirical brain networks^40^. In contrast, lazily trained networks maintained a Fisher kurtosis of 0 (weight kurtosis change, Rich=0.27, training=0.001, Rich vs. lazy t_18_=192.20, p<10e-31) (Fig. 7i). Finally, we characterized the dimensionality of the weights to gain insight into the successive representational transformations in the ANN. Specifically, we performed singular value decomposition (SVD) on the weights and calculated the participation ratio of the singular values. We found that that weight dimensionality was lower for rich vs. lazy training regimes (participation ratio: 142.31 vs. 47.92; t_18_=-913.86, p<10e-44) (Fig. 7j). These findings suggest that across layers, richly trained ANNs with low-dimensional weights collectively produced highly diverse and modular patterns of representations across layers, consistent with empirical data (Fig. 7c,d).

### Transformational trajectories from visual to motor representations

To provide a state-space intuition of the different representational transformations in rich and lazy ANNs, we characterized the transformational trajectories from visual to motor representations. This involved plotting ANN transformations in a 2D space. The y-axis reflected the alignment (inner product) with visual representational geometry and the x-axis reflected the alignment with motor representational geometry. While a single linear transformation would map directly visual to motor representations (dashed line; Fig. 8a), we hypothesized that compression-then-expansion would occur by first compressing representations along the visual axis, followed by expansion along the motor axis (blue curve, Fig. 8a). This is in contrast to the alternative, where motor representations first expand with minimal loss of visual representations (red curve in Fig. 8a). In agreement with this theory, we found that richly trained ANNs first compressed along the visual axis, followed by growth along the motor axis (Fig. 8b). This is consistent with the notion that higher-order brain areas (i.e., similar to intermediate layers in an ANN) contain functionally distinct representations from input (visual) and output (motor) representations, and are low-dimensional (Fig. 6c). In contrast, lazy ANN representations maintained high similarity to visual input representations, with visual-to-motor representational transformations primarily implemented in the final read-out weights.

**Figure 8.**
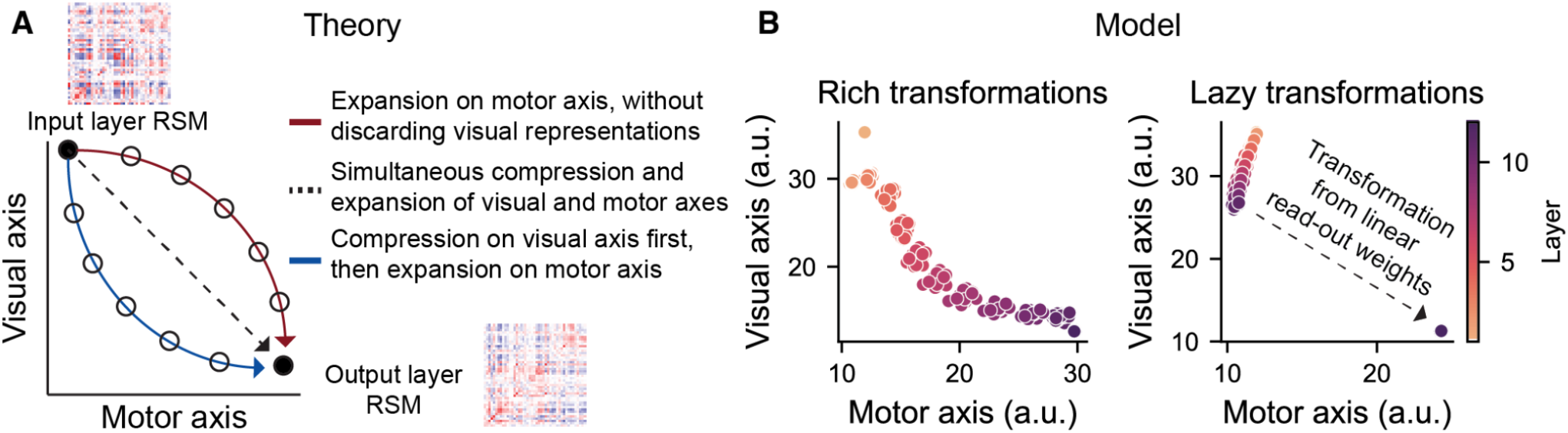
Trajectories of representational transformations from visual to motor content. **a)** A theory of how representations transform across layers/brain areas, from visual to motor representations. Axes reflect the similarity (computed as the inner product) to the visual input region’s RSM (y-axis) and the motor output region’s RSM (x-axis). Hidden layer representations can then be plotted along these two dimensions by calculating the inner product between the sensory and motor RSMs. **b)** We plotted the ANN’s internal representations along these two dimensions, finding that rich representations are consistent with compression first along the visual axis, then expansion along the motor axis. In contrast, lazy ANNs preserve visual representations in hidden layers until the final read-out weights transform visual into motor representations. Note that y- and x-axis are not necessarily orthogonal, and are plotted as such for visualization purposes. Each dot in the scatter plots reflects a different ANN initialization and layer.

## Discussion

Using RSA-based techniques, we mapped the multi-task representational organization of human cortex. We found that RA segregation (an inter-region measure) and the representational dimensionality (a regional measure) was associated with the unimodal-transmodal hierarchy. However, when evaluating the dominant axis of variation of cortico-cortical RA, we found that the principal RA gradient was organized along a sensory-association-motor axis. This revealed that representations compressed-then-expanded from sensory perception, to integrative representations in association areas, to motor action. To explore the computational mechanisms of hierarchical multi-task representations in the brain, we used feedforward ANNs to study how representations compressed-then-expanded from input to output. We found that during a rich ANN training regime, the ANN learned structured and hierarchical representations that 1) compressed-then-expanded representations and 2) had greater similarity to representations found in fMRI data. Further analysis of the ANN revealed that this training regime produced low-dimensional connectivity weights with a heavy-tailed distribution, consistent with observations made in empirical brain networks^40^. Together, these findings characterize the topographic organization of multi-task representations in cortex, and provide a framework for understanding how these representations emerge in computational models.

Meta-analytic studies that aggregate data across experiments have previously provided coarse-grained atlases of cognition across the brain^9,10^. However, because aggregating data across subjects, experimental designs, and MRI scanning protocols can make it difficult to directly compare tasks directly, these studies primarily mapped the univariate activations of individual brain regions to tasks. One recent study collected many tasks per subject and, by using individualized encoding models, identified the clustering of tasks in a latent cognitive space^41^. Our study complements that study by focusing instead on the cortical topography of multi-task representations, rather than relationships of tasks in a latent encoding space. The generalizability of findings from any multi-task study is limited by the selection of tasks included in the experimental design. Thus, it will be important for future studies to evaluate other task types and datasets, including modular and ecologically valid tasks^42^. Nevertheless, the insights gained from multi-task studies illustrate the importance of collecting significant amounts of data *per subject* to reveal fine-grained relationships between neural and cognitive processes^17,43,44^.

Our findings of hierarchical gradients in task representations adds to a growing literature identifying hierarchy as a fundamental principle of cortical organization. Early seminal work using tract-tracing techniques revealed hierarchical connectivity organization in the macaque visual cortex^45^. More recent work has shown that such hierarchical organization can be studied in humans *in vivo* with structural and functional MRI and electrophysiology technology. These studies have focused on identifying structural^32^, transcriptomic^31^, RSFC^30^, and intrinsic timescale signatures of hierarchical organization^46,47^. Most of these hierarchical descriptions used task-free MRI data, and here we establish an overarching link that bridges multi-task representations with fundamental hierarchical organization. Other studies that evaluate the role of functional and anatomical connectivity organization have also identified network hubs in association regions^48,49^. This is consistent with our finding that association areas contain integrative representations that link the sensory and motor representations that lie on opposing ends of the sensory-motor axis. Future studies can explore how specializations of association cortex, in long-range anatomical connectivity and local microcircuitry, contribute to the formation of low-dimensional integrative representations.

We found that a rich training regime produced hidden representations that were significantly more similar to empirical brain representations, suggesting that the brain also learns feature-rich representations. What is the utility of low-dimensional, feature-rich representations in the brain? One recent study, by Flesch and colleagues, suggested that feature-rich representations are useful for transfer learning and generalization to novel conditions^21^. Specifically, they showed that feature-rich learning projects input features onto low-dimensional, orthogonal manifolds that minimize inference while maximizing the robustness of task information, producing generalizable representations. Consistent with their findings generalizable representations in frontoparietal areas, here we found low-dimensional representations across association cortex. Low-dimensional representational geometries are likely useful for generalization, in brains and in models, because many distinct tasks recruit shared neurocomputational mechanisms, such as modular processing. Moreover, recent work suggests that learning multiple different tasks naturally produces low-dimensional and abstract representations^50^. Thus, it will be important for future studies to provide a unified understanding of the contribution of low-dimensional representations for rapid reconfiguration and generalization to novel tasks^51^.

Our computational modeling results provide a parsimonious framework to study representational transformations in relation to empirical data. There are multiple directions in modeling and analytics that future studies can explore. First, we used a simple feedforward ANN, motivated by our findings of a dominant sensory-to-motor gradient. Future models can examine the impact of more complex and recurrent ANN architectures of internal representations^52,53^. We found that representations depended strongly on the training regime, which we controlled by weight initialization following prior literature^21,22^. Future modeling should explore alternative training methods for ANNs to examine how they alter the similarity to empirically observed representations. For instance, training strategies, such as continual and/or curriculum learning, can promote modularity of internal representations and generalization of task performance^20,54^. Finally, future studies should examine the metrics used to quantify structure and similarity in empirical and model representations. For instance, inherent constraints on RSMs can be used to define alternative measures of RA^55,56^. Therefore, it will be important for future work to explore the space of biologically-relevant strategies that produce feature-rich, hierarchical representations in models which can be quantitatively related to neural datasets.

In conclusion, we characterized the geometry and topography of 26 diverse cognitive tasks across the human cortical hierarchy, and provide insight into the computational mechanisms that produce similar representations in ANNs. Overall, analysis of the task representational hierarchy revealed a sensory-to-motor gradient that compressed-then-expanded task representations. Subsequent modeling of these task activations in ANNs revealed that a rich training regime can reproduce representations that were consistent with empirical data. This finding provides a framework to explore how to build ANNs that learn task representations in a brain-like manner. We expect these findings to spur new investigations into how the study of multi-task representations in the brain can inform new models of multi-task performance in machine learning models.

## Methods

### Multi-domain task battery dataset

Portions of this section are paraphrased from the dataset’s original publication’s Methods section^24^. We used the publicly available multi-domain task battery (MDTB) dataset^24^. The MDTB dataset contains task fMRI data for 24 subjects collected at Western University (16 women, 8 men; mean age = 23.8 years, s.d. = 2.6; see^24^ for exclusion criteria). All participants self-reported to be right-handed. Briefly, the MDTB dataset contains 26 unique cognitive tasks with up to 45 different task conditions per participant. Participants first scanned all tasks in set A, and returned for a second session to perform tasks in set B (Fig. 2a). Each task set session consisted of two imaging runs each. Half of the subjects had sessions separated by 2-3 weeks, while the other half had sessions separated by a year. Of the 24 subjects, a separate resting-state fMRI scan was collected for 18 subjects. Resting-state FC analyses presented in Fig. 3 were performed using this subset of subjects.

A large battery of tasks was selected to broadly recruit cognitive processes from many functional domains (Fig. 2a). Set A consisted of cognitive, motor, affective, and social tasks. Set B contained eight tasks that were also included in set A (e.g., theory of mind and motor sequence tasks), and nine unique tasks. Both sets contained 17 tasks each. Additional details regarding the experimental tasks and conditions have been previously reported (see Supplementary Table 1; https://static-content.springer.com/esm/art%3A10.1038%2Fs41593-019-0436-x/MediaObjects/41593_2019_436_MOESM1_ESM.pdf)^24^.

Tasks were performed once per imaging session for 35 s blocks. Task blocks began with a 5 s instruction screen followed by 30 s of continuous task performance. While most tasks consisted of 10-15 trials per block, the number of trials per task ranged from 1 to 30 (e.g., go/no-go task versus movie watching). 11 of the 26 tasks were passive, meaning no behavioral responses were required (e.g., movie watching). For the remaining tasks, responses were made with either left, right, or both hands using a four-button box. Responses were made with either index or middle fingers of the assigned hand(s). Performing all tasks within a single imaging run for each participant ensured a common baseline between tasks, enabling fine-grained multi-task analyses. Additional details regarding trial structure, inter-trial-interval timings, etc., have been previously reported^24^

### fMRI preprocessing

Resting-state and task-state fMRI data were minimally preprocessed using the Human Connectome Project (HCP) preprocessing pipeline within the Quantitative Neuroimaging Environment & Toolbox (QuNex)^57,58^. The HCP preprocessing pipeline consisted of anatomical reconstruction and segmentation, EPI reconstruction and segmentation, spatial normalization to the MNI152 template, and motion correction. Additional nuisance regression was performed on the minimally preprocessed time series. Consistent with previous reports^59^, this included six motion parameters, their derivatives, and the quadratics of those parameters (24 motion regressors in total). We also removed the mean physiological time series extracted from the white matter and ventricle voxels. We also included the quadratic, derivatives, and the derivatives of the quadratic time series of each of the white matter and ventricle time series (8 physiological nuisance signals). This amounted to 32 nuisance parameters in total, and was a nuisance regression model that was previously benchmarked^60^. In addition to nuisance regressors, task fMRI data was also modeled with task regressors to extract activation estimates described below.

### fMRI task activation estimation

We performed a single-subject task GLM analysis on fMRI task data to estimate vertex-wise surface activations for each task condition on the CIFTI grayordinate space^61^. We modeled a separate regressor for every trial within each imaging run, similar to a beta series model (Rissman et al., 2004). The instruction period for each task was not included in the task regressors. This enabled the estimation of specific task conditions within each task block (e.g., congruent versus incongruent conditions for the Stroop task). Each regressor (trial) was modeled as a boxcar function from the onset to offset of the trial (0s indicate off, 1s indicate on), and then convolved with the SPM canonical hemodynamic response function to account for hemodynamic lags^63^. Activations for a task condition were then obtained by averaging the activation beta coefficients across trials within each imaging run, resulting in one task condition activation per run. Task GLMs were performed using the LinearRegression function within scikit-learn (version 0.23.2) in Python (version 3.8.5).

### Representational similarity analysis and representational alignment

We performed a split-half, cross-validated RSA to characterize the geometry of task representations across cortex (Kriegeskorte et al., 2008). RSA was performed for each parcel in the Glasser et al. (2016) atlas using vertices within each parcel^23^. Critically, RSA was performed at the subject-level to ensure that fine-grained representations were subject-specific and that activations would not be averaged across subjects. (Group averaging was computed after RSMs were constructed for each subject at every parcel.) We used all task conditions, resulting in a 45 × 45 RSM. We used cosine similarity to measure the distances between task activations. Despite many alternative metrics^64,65^, we specifically chose the cosine similarity, since it also takes into account the overall mean magnitude of activation across a set of vertices (in contrast to Pearson correlation). Cross-validation was achieved by measuring the cosine similarity of activation patterns of the first and second imaging sessions (i.e., a split-half cross-validation). This was possible since all tasks (in set A and B) were performed in two separate imaging runs. This ensured a non-trivial diagonal element (i.e., not equal to 1), which revealed the test-retest reliability (or similarity) of the activation patterns of the same task condition.

Inter-regional representational alignment (RA) was calculated by measuring the cosine similarity of the upper triangle elements (including the diagonal) of two region’s RSMs. Related measures have also been previously introduced under the term “representational connectivity”^4,55,66^.

### Network segregation

Network segregation for RSFC and multi-task RA was measured as the difference between within-network and between network FC/RA, divided by within-network FC/RA^29^. Networks were defined using a previously published whole-brain resting-state network partition^27^. Network segregation^29^ was calculated for each region separately using either the RA or FC matrix. Specifically, the segregation *S_region_* of a region was calculated as

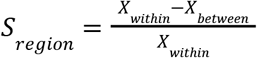

where *X_within_* is the within-network FC/RA for the region of interest and *X_between_* is the out-of-network FC/RA.

### Representational dimensionality and multi-task decoding

Representational dimensionality was measured as the participation ratio of the multi-task RSM. The participation ratio was calculated as

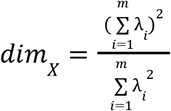

where *dim_X_* corresponds to the representational dimensionality of region *X*, and λ_*i*_ corresponds to the eigenvalues of the RSM of region *X* with *m* eigenvalues. Intuitively, the flatter the eigenspectrum of region *X*’s RSM, the higher the dimensionality.

To complement representational dimensionality, we also measured the multi-task decodability (45-way classification) of each region using a minimum-distance classifier. We used the cosine angle as our measure of distance and split-half cross validation. Thus, a successful classification indicated that the diagonal element of a region’s cross-validated RSM was greater (i.e., smallest distance) than all other off-diagonal elements for a given row (see Fig. 4a).

We performed additional control analyses to account for parcel size (i.e., the number of vertices) when calculating representational dimensionality and multi-task decodability. This was performed by conditioning on (regressing out) the number of vertices from each measure using linear regression (regression was performed across parcels). We then re-calculated the correlation across brain maps (e.g., myelin map vs. representational dimensionality) using the residual values (Supplementary Fig. 3b,c).

### Gradient analysis

Cortical gradients were calculated using a principal component analysis (PCA) on parcellated data. For resting-state FC gradients, PCA was applied on the cortical FC matrix. For RA gradients, PCA was applied on the cortical RA matrix. Consistent with previous studies (Margulies et al., 2016), matrices were thresholded to include only the top 20% of values prior to extracting gradients. All correlation-based statistical tests involving gradients (i.e., spatial correlations across cortex) were performed using spatial autocorrelation-preserving permutation tests that generated random surrogate brain maps (Burt et al., 2020). We used the BrainSMASH toolbox to generate 1000 random surrogate brain maps for each cortical map of interest, and non-parametric p-values were calculated from the null distribution. Therefore, the lowest precision non-parametric p-value we obtained was 0.001.

### Testing for compression-then-expansion in empirical data

Assessing compression-then-expansion in empirical data involved fitting representational dimensionality to sensory-motor hierarchy loadings using regression models (Fig. 5d and Supplementary Fig. 5). We specifically used RSFC sensory-motor gradient 2 loadings (*x* variable) as the regressor to predict the representational dimensionality of each parcel (*y* variable). For model adjudication, we used used several competing regression models, including:

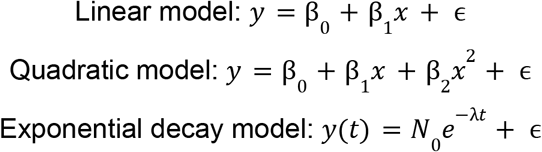

where β_*i*_ was the fitted coefficient term, and ϵ was the residual error term. For the 2nd-order quadratic model, a positive 2nd-order coefficient indicated a convex quadratic. Selection of the model was based on the lowest Akaike Information Criterion and Bayesian Information Criterion (Supplementary Fig. 5).

To further verify compression-then-expansion across the sensory-motor hierarchy, we binned together groups of 10 bins of 36 parcels according to their RA principal gradient loading (Fig. 5a). To establish compression-then-expansion along this gradient, we fit a piecewise linear model with the functional form

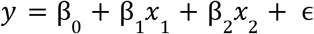

We trained a piecewise linear model for every possible breakpoint (i.e., where *x*_1_ < *i* and *x*_2_ > *i* for every bin *i* between 1 and 10; 8 possible models). Note that *x*_1_ were values for *x* < *i* and 0 otherwise, and *x*_2_ were values for *x* > *i* and 0 otherwise. After identifying the model with the greatest fit evaluated using R^2^, which turned out to be the model with the breakpoint at *i* = 3, we tested the statistical significance for the beta coefficients β_1_ and β_2_, with the hypothesis that they should be negative and positive, respectively. A negative and positive slope for β_1_ and β_2_ would reflect a compression of representational dimensionality from input to the breakpoint, and then an expansion from the breakpoint to the output.

### ANN modeling and training

We modeled the transformation from visual fMRI activations to motor activations using a linear feedforward ANN. This enabled the characterization of the transformation as a sequence of linear transformations. fMRI activations were selected based on lying on opposite ends of the RSFC sensorimotor gradient (i.e., region with the lowest/highest loadings). Input activations were normalized across vertices before training. Inputs and outputs corresponded to the vertex-level fMRI task activations for each parcel. We used the RSFC sensorimotor gradient rather than the task-based RA gradient to avoid any potential confounds of selecting activations from the same task data. The input and output parcels corresponded to parcel 338 and 235 in the Glasser et al. (2016) atlas, respectively (Glasser et al., 2016). We built the ANN with 10 hidden layers with tied weights (500 units per layer), and was defined by the equations

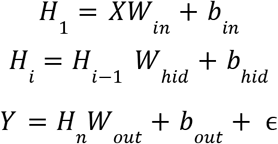

where *X* was the input fMRI activation from the visual parcel, *W_in_* mapped vertex activations into the hidden unit space, *b_in_* were the input biases, *H_i_* was the hidden unit activations for layer *i*, up to *n* (i.e., 10), *W_hid_* and *b_hid_* were the weights and biases for the hidden layers, *Y* was the predicted motor fMRI activation in the motor parcel, and ϵ was the residual error term. Using tied weights and a linear model reduced the number of free parameters in the model, thereby constraining the solutions and simplifying the model for subsequent analysis. Using tied weights also increased computational efficiency during training. However, we also ran the model without tied weights (where *W_hid_* and *b_hid_* were distinct for each layer), yielding computationally similar results (Supplementary Fig. 6).

ANN hidden layer weights were initialized from a Xavier normal distribution, with mean 0 and a scaling factor ranging from 0.2 to 2.0 in increments of 0.2^68^. Biases were initialized to be 0. Training was implemented using a mean squared error cost function and the Adam optimizer with an initial learning rate of 0.0001^69^. Training was stopped once the mean squared error fell below a threshold of 0.2.

We fit ANNs for each subject’s activations separately. For every subject, we trained 20 networks with different random initializations. For each ANN analysis, statistics and network properties (e.g., dimensionality, weight norms, etc.) were averaged across subjects, and statistical tests were performed on the 20 random initializations.

All models were built using PyTorch version 1.4.0 and Python version 3.8.5.

### ANN analysis

Trained ANNs were subject to analysis to characterize both the learned intermediate representations and weight distribution properties. Model RSMs were generated by propagating subject-level activations across all tasks through the hidden layers. Cross-validated RSMs were constructed and analyzed identically to fMRI data (e.g., cosine similarity and then participation ratio to estimate its dimensionality; Fig. 6c). As in our fMRI analysis, we used a split-half cross-validation where we compared task activations between the first and second imaging sessions of each task set. We fitted the dimensionality across ANN layer depth using a 2nd-order polynomial regression to assess how representational dimensionality changed throughout the network (Fig. 6d). A positive and negative 2nd-order coefficient indicated a convex and concave quadratic, respectively.

We compared the representational geometries produced by the ANN with the representational geometries found in empirical fMRI data. To directly compare ANN and empirical RSMs, we partitioned cortex into 10 bins containing 36 parcels each (Fig. 6g). Cortical bins and their ordering were determined by the RSFC sensory-motor gradient, where parcels with similar loadings (were placed in adjacent bins) (Fig. 5a). We computed the cosine similarity of each region’s RSM with each ANNs layer’s RSM. To evaluate the correspondence between representations in each cortical bin and each ANN layer, we averaged the cosine values across parcels within each bin (Fig. 6f). This was done for ANNs trained under the rich regime (weight initializations less than 1) and the lazy regime (weight initializations greater than 1).

We assessed the inter-layer RA within the ANN for different weight initializations (Fig. 7a), which is similar to inter-region RA measured in fMRI data (Fig. 3c). This was defined as the cosine similarity between RSMs between pairs of ANN layers (Fig. 7a). We also analyzed the properties of the trained and initialized ANN weights. This included calculation of the Frobenius norm, Fisher kurtosis, and singular value decomposition of the weight matrices under different weight initializations. Dimensionality of the ANN’s weights were performed by measuring the participation ratio of the singular values. All statistical analyses were carried out in Python version 3.8.5 using the NumPy (version 1.18.5) and SciPy (version 1.6.0) packages.

## Data availability

All data in this study has been made publicly available on OpenNeuro by King and colleagues^24^. (URL: https://openneuro.org/datasets/ds002105/)

## Code availability

All code related to this study will be made publicly available on GitHub. Analyses and models were implemented using Python (version 3.8.5).

## Acknowledgements

This project was supported by NIH grant R01MH112746 (JDM) and a Swartz Foundation Fellowship (TI). The authors acknowledge the Yale Center for Research Computing at Yale University for providing access to the Grace cluster and associated research computing resources. We thank Warren Pettine, Markus Helmer, and Jacob Miller for comments on earlier drafts of the manuscript.

## Supplementary Figures

**Supplementary Figure 1.**
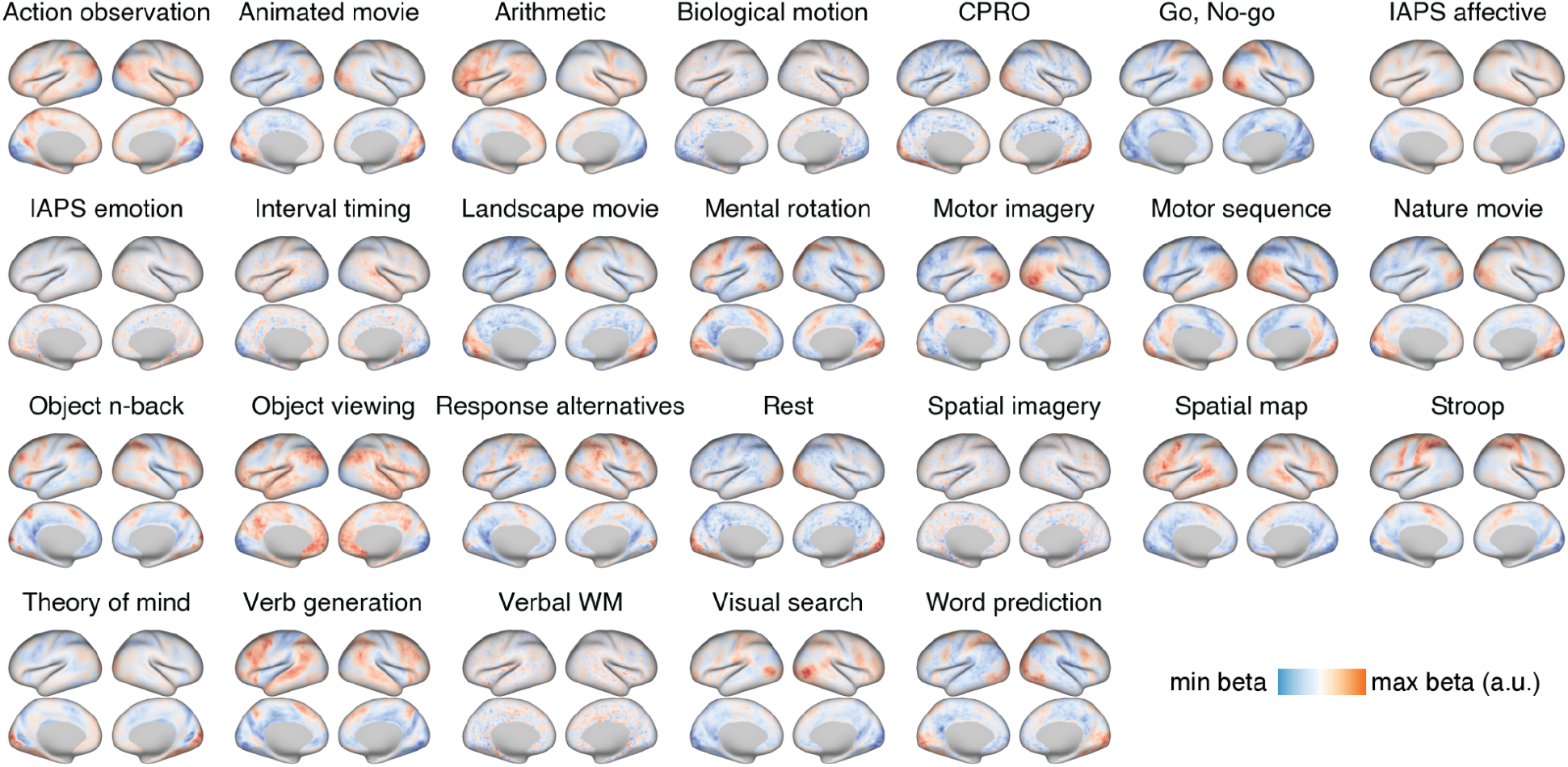
Whole-cortex group activation maps for all 26 cognitive tasks. Activation maps reflect the GLM beta values, and were averaged across conditions within each task.

**Supplementary Figure 2.**
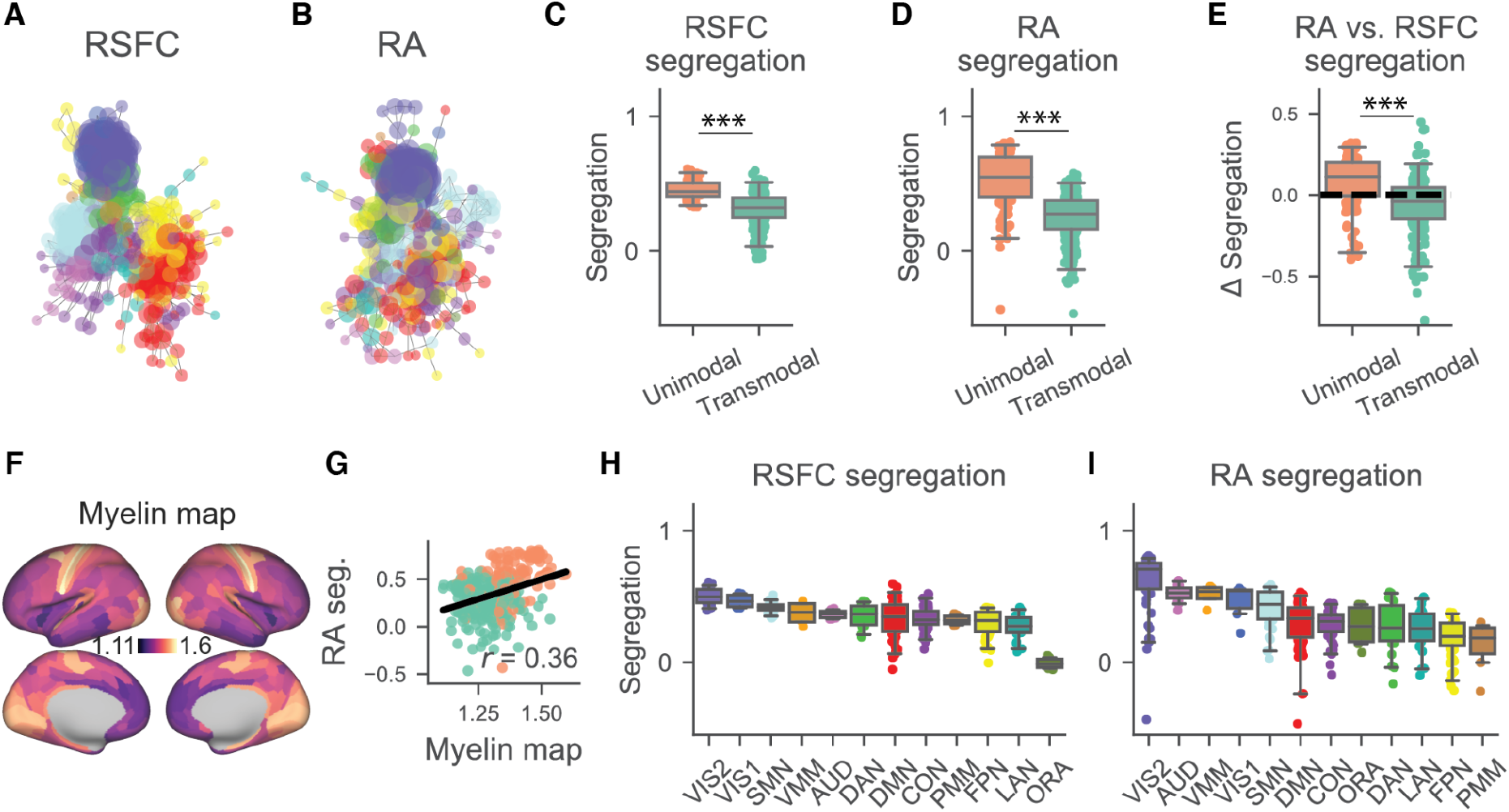
Comparing segregation of whole-cortex RSFC and RA between unimodal-transmodal areas and functional networks. **a, b)** Force-directed graphs comparing RSFC and RA community structure (color-coated by functional networks). **c)** Segregation of RSFC and **d)** RA whole-cortex matrices. **e)** The direct comparison of differences in segregation between RA and RSFC for unimodal and transmodal regions (same as Fig. 3h). **f, g)** Association of regional RA segregation with the cortical myelin map (T1w/T2w structural map). **h)** Segregation of RSFC by functional networks. **i)** Segregation of RA by functional networks. Note that for both RA and RSFC, sensorimotor networks have higher segregation than association networks. Network key: VIS1=Visual 1; VIS2=Visual 2; SMN=Somatomotor; VMM=Ventral multimodal; AUD=Auditory; DAN=Dorsal attention; DMN=Default mode; CON=Cingulo-opercular; PMM=Posterior multimodal; FPN=Frontoparietal; LAN=Language; ORA=Orbital-affective. Colors of each network correspond to colors in panel Fig. 3e)

**Supplementary Figure 3.**
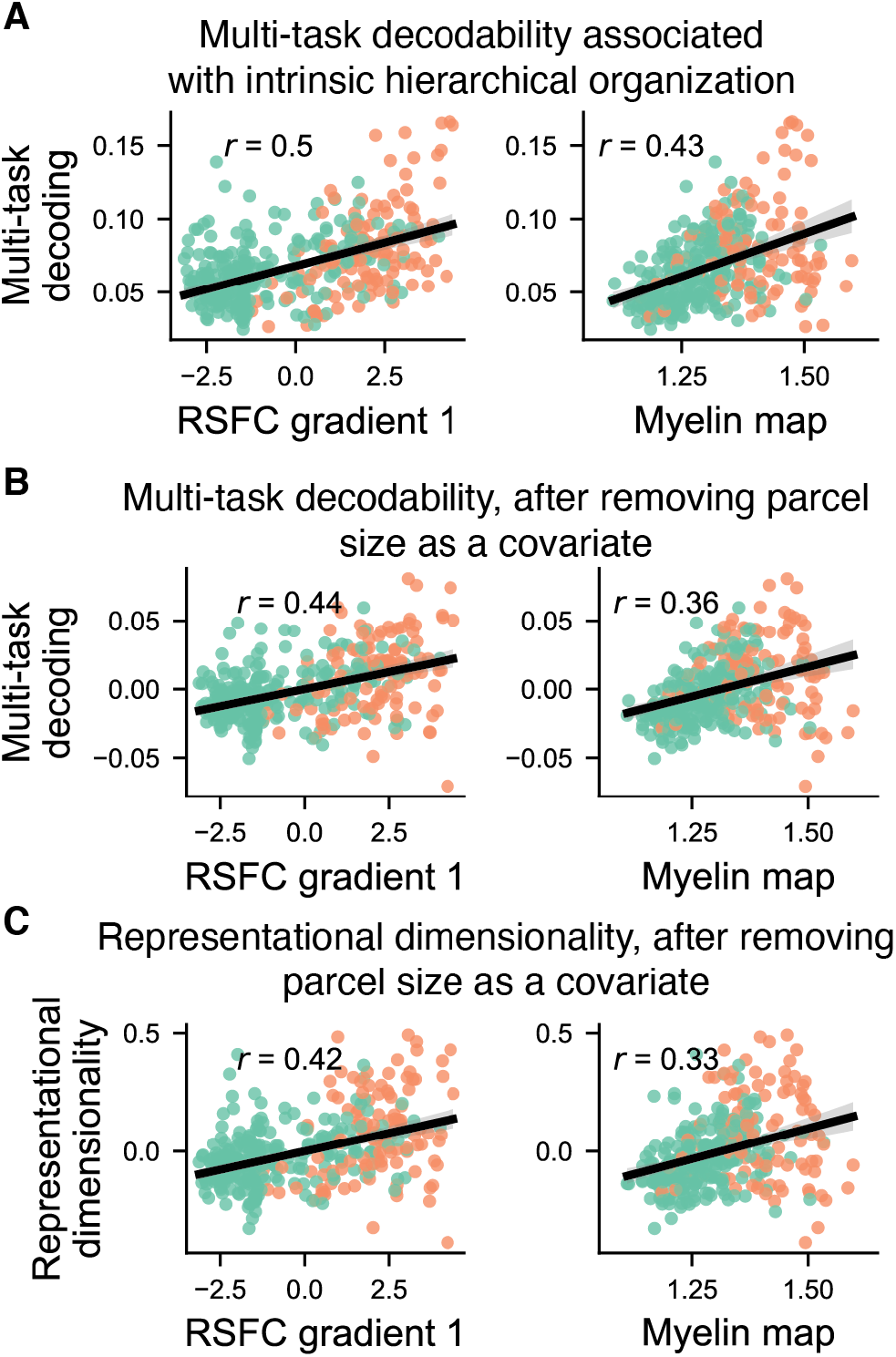
Representational dimensionality and multi-task decoding produce similar associations with intrinsic hierarchy, even after controlling for parcel size. **a)** Correlation of multi-task decoding with the principal RSFC gradient and myelin map across regions. **b)** After removing parcel size (i.e., the number of vertices within each parcel) as a covariate (via linear regression), a strong association between decodability and intrinsic hierarchy was maintained. **c)** Same analysis as in panel **b**, but using representational dimensionality rather than decodability. All correlations in a, b, and c resulted in a non-parametric p<0.001 using surrogate brain maps that accounted for spatial autocorrelation^67^

**Supplementary Figure 4.**
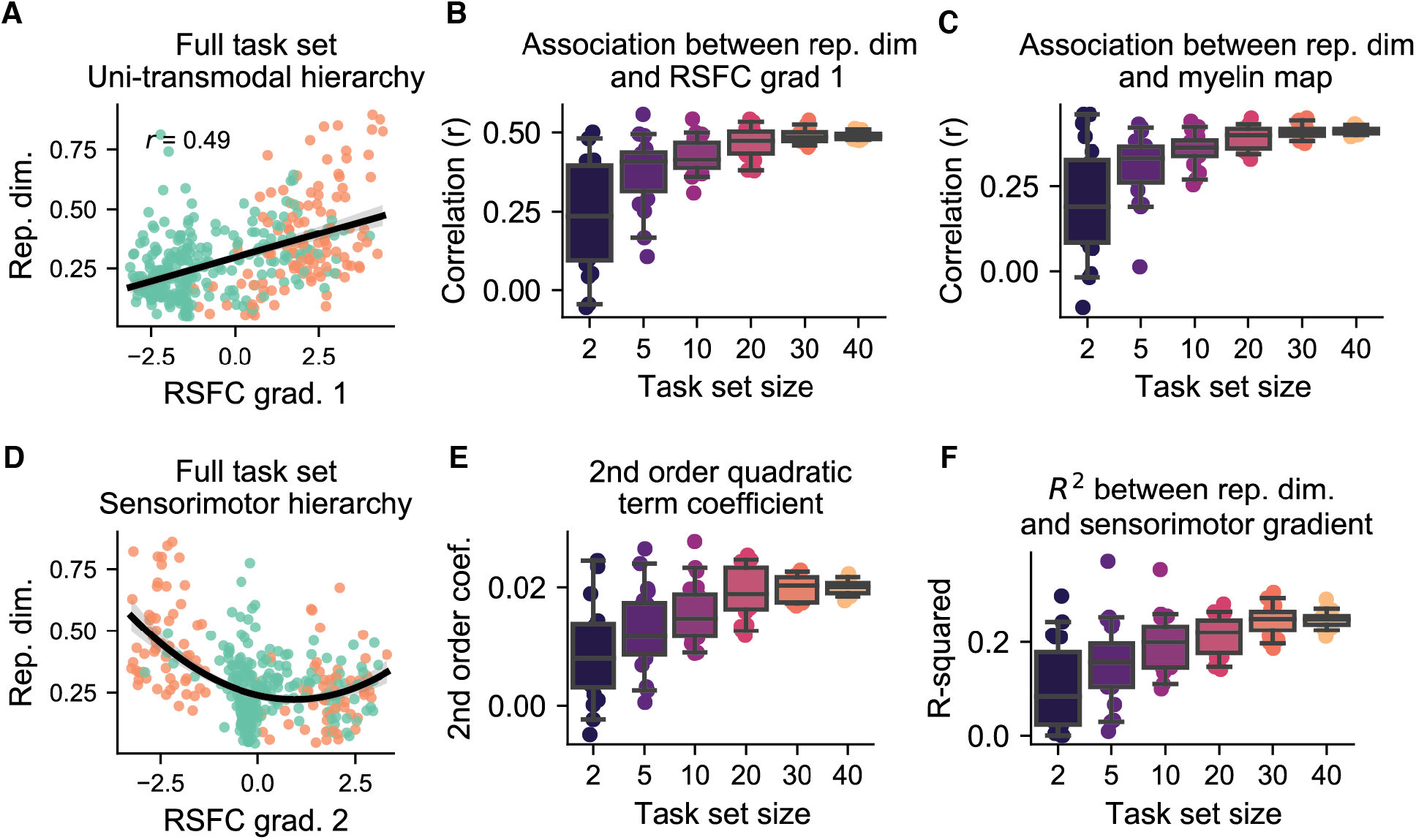
Random subsamples of the task set show similar association with both the unimodal-transmodal and the sensorimotor hierarchy. **a)** The association between representational dimensionality and the principal RSFC gradient (unimodal-transmodal hierarchy) with the entire task set. **b)** We randomly sub-sampled (without replacement) tasks to downsize the RSMs of all parcels, and then measured the correlation between representational dimensionality and RSFC gradient 1. For each sub-sample size, we repeatedly chose (i.e., 45 choose n) 20 times to estimate the robustness of the association with arbitrary selection of tasks. We found that the association increased and stabilized as we increased the number of tasks. **c)** Same as in **b**, but using the myelin map. **d)** The compression-then-expansion fit of representational dimensionality and the sensorimotor (RSFC gradient 2) hierarchy. We estimated the 2nd-order polynomial fit for randomly sub-sampled tasks, and assessed the coefficient of 2nd-order polynomial fit. The higher (and more positive) the parameter, the more convex the compression-then-expansion was. We found increased compression-then-expansion as the number of randomly sampled tasks were included. **f)** Same procedure as **e**, but measuring the R-squared of the polynomial fit rather than the 2nd-order coefficient term.

**Supplementary Figure 5.**
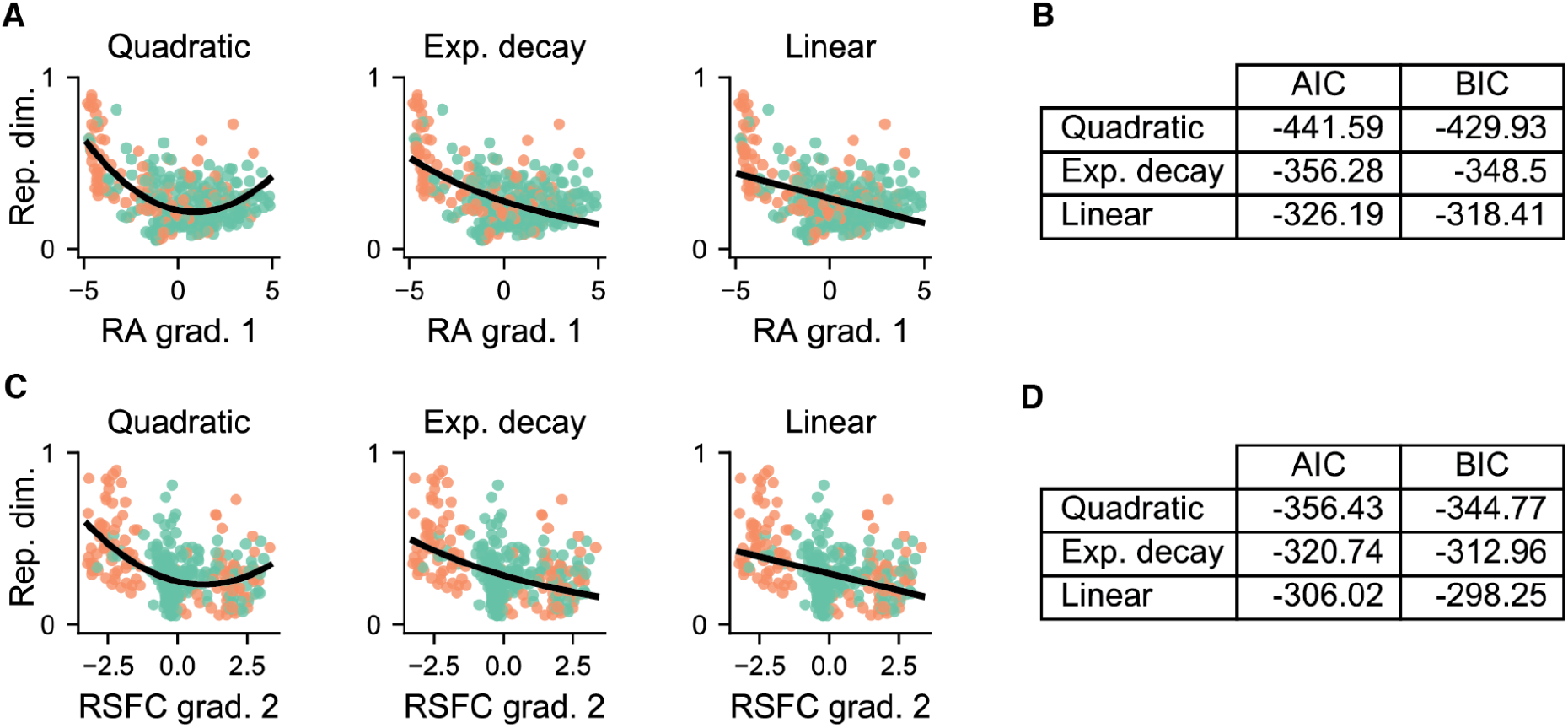
Establishing compression-then-expansion of representational dimensionality across the sensory-motor hierarchy via model adjudication. **a)** We fit the representational dimensionality of parcels across the sensory-motor RSFC gradient using three competing models: Quadratic (2nd-order polynomial), linear, and an exponential decay model, where separate models were fit for loadings less than and greater than 0. **b)** The Akaike Information Criterion (AIC) and the Bayesian Information Criterion (BIC) for all models, which takes into account the maximum likelihood of each model while penalizing the number of parameters. Quadratic models had the smallest values for both AIC and BIC. **c,d)** Same as panels **a** and **b**, but using the RA principal gradient. Quadratic models were defined as *y* = β_0_ + β_1_*x* + β_2_*x*^2^ + ϵ. Linear models were defined as *y* = β_0_ + β_1_*x* + ϵ. Exponential decay models were defined as *y*(*t*) = *N*_0_*e*^−*λt*^ + ϵ.

**Supplementary Figure 6.**
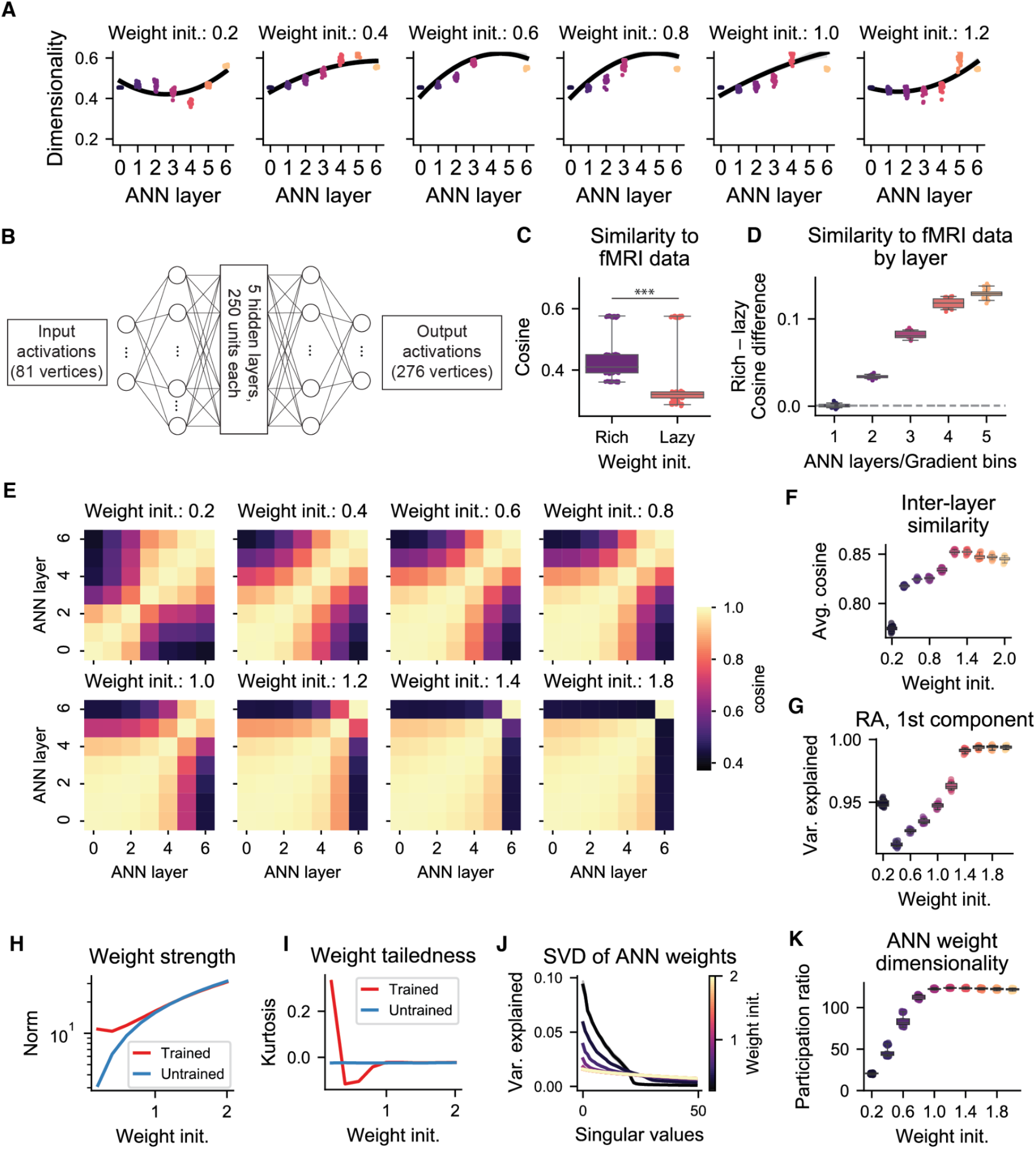
Training an ANN with untied weights results in qualitatively similar results. We trained a 5-layer ANN with untied weights to produce qualitatively similar results to the ANN in the main manuscript. We reduced the number of layers from 10 to 5 and the number of hidden units from 500 to 250 for computational efficiency. (An ANN with untied weights has significantly greater parameters than one with tied weights.) **a)** Representational dimensionality of ANN layers for different weight initializations. **b)** ANN architecture. **c)** Richly trained ANNs had significantly higher similarity with representations found in empirical data relative to lazily trained ANNs. **d)** Similarity to fMRI data by layer (rich minus lazy ANNs). **e)** Representational alignment of each ANN’s layer (cosine similarity between RSMs). **f)** Overall similarity of representations across ANN layers. Greater representational dissimilarity (across layers) is found in richly trained ANNs. **g)** Variance explained of the first principal component for each of the RA matrices in panel **e**. **h)** Frobenius norm of the weight distribution across initializations. **i)** The kurtosis (tailedness) of the weight distribution across layers under different weight initialization schemes. **j)** SVD of ANN weights. **k)** Dimensionality (participation ratio) of the weights for different initializations. Richer training regimes produce low-dimensional weights.

**Supplementary Figure 7.**
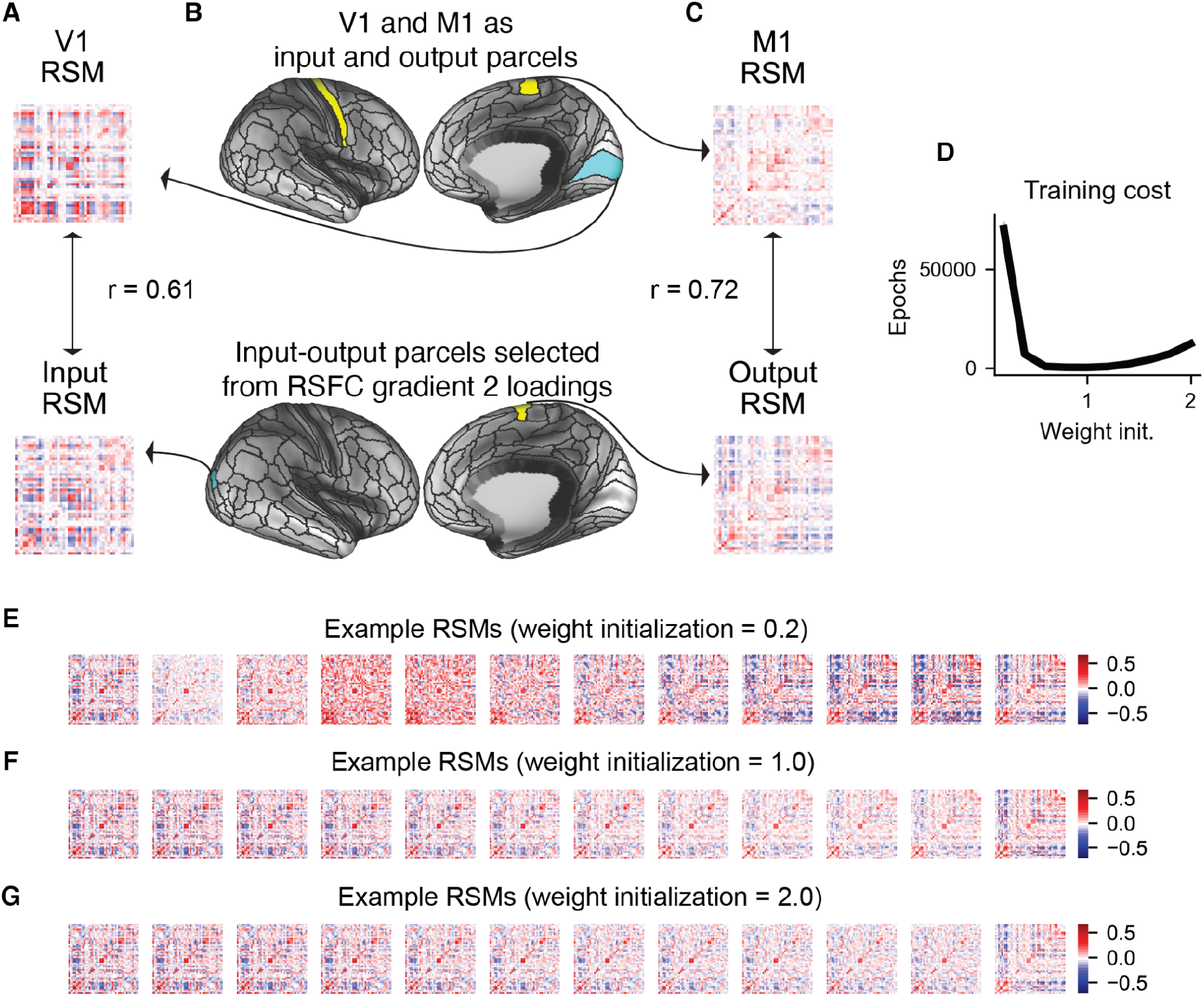
Supplemental information ANN modeling during rich and lazy training regimes. The similarity between **a)** the RSMs for V1 and the gradient-identified input parcel for model construction and **b)** the RSMs for M1 and the gradient-selected motor output parcel. Overall, the representational geometries were highly similar between V1 and the input RSM, and M1 and the motor output RSM. **d)** The training cost (i.e., number of training epochs required) for different weight initializations. Visualization of RSMs for example ANNs (one initialization each) for **e)** rich, **f)** intermediate (i.e., initialization SD=1.0), and **g)** lazy training regimes.

